# Gelsolin Counteracts ER Stress-Driven Inflammatory Circuits in Psoriasis-like Dermatitis

**DOI:** 10.1101/2025.11.20.689413

**Authors:** Daisuke Ori, Haruna Okude, Riko Konishi, Motoya Murase, Shuya Hiroki, Saki Takahara, Towa Tanaka, Rina Toyodome, Norisuke Kano, Takumi Kawasaki, Ken J Ishii, Kouji Kobiyama, Hideyuki Nakashima, Kinichi Nakashima, Miwa Sasai, Masahiro Yamamoto, Yutaro Kumagai, Akio Tsuru, Kenji Kohno, Taro Kawai

## Abstract

Psoriasis is a chronic inflammatory skin disorder driven by amplified communication between immune cells and keratinocytes. Here, we show that imiquimod (IMQ) triggers organelle stress responses that directly contribute to this pathogenic circuit. In dendritic cells (DCs), IMQ promotes formation of ER–mitochondria contact sites (MAMs), inducing ER stress and activation of the unfolded protein response (UPR). These pathways act independently of, yet converge with, TLR7/MyD88 signaling to enhance IL-23 expression. IMQ also increases cytosolic Ca²^+^, facilitating NLRP3 inflammasome activation and release of mitochondrial DNA (mtDNA). In parallel, keratinocytes exposed to IMQ activate UPR-dependent genes, including Defb14 (mBD14), a psoriasis-associated antimicrobial peptide. Extracellular mtDNA and mBD14 then cooperatively stimulate plasmacytoid DCs through TLR9, establishing a feed-forward inflammatory loop. We further identify Gelsolin as a direct IMQ-binding protein that mitigates IMQ-induced ER stress; its loss amplifies ER stress, UPR activation, and oxidative stress, and its expression is reduced in human psoriatic lesions. Thus, MAM–UPR signaling links intracellular organelle stress to the intercellular networks that drive psoriatic inflammation, with Gelsolin acting as a critical intrinsic safeguard.

## Introduction

Psoriasis is a chronic inflammatory skin disease characterized by excessive keratinocyte proliferation and abnormal differentiation, and immune cell infiltration into the epidermis^1,2^. The interplay between keratinocytes and immune cells, including dendritic cells (DCs) and T cells, drives disease pathogenesis, with the IL-23/IL-17A axis playing a central role^3,4^. IL-23, primarily secreted by DCs, promotes the expansion and maintenance of IL-17-producing T-cell populations, particularly Th17 and Tc17 cells, which are considered major sources of IL-17A in human psoriasis^3,5–8^. Although IL-17- producing γδT cells play an important role in murine psoriasis-like models, their contribution to human psoriasis is less prominent^9^. IL-17, in turn, binds keratinocyte receptors, triggering epidermal hyperproliferation and the release of chemokines and antimicrobial peptides (AMPs)^4,10,11^. These factors recruit immune cells to psoriatic lesions, amplifying IL-23 production and establishing a self-perpetuating inflammatory loop^3,11,12^. FDA-approved biologics, such as secukinumab, ixekizumab, and guselkumab, target IL-23 or IL-17A to mitigate symptoms^3,13^. Additionally, IL-1β from DCs enhances Th17 activation^14,15^, while plasmacytoid DCs (pDCs) infiltrate lesions and secrete type I interferons and proinflammatory cytokines^16^. Notably, AMPs like LL37 form complexes with self-DNA/RNA from damaged cells, activating pDCs via TLR7 or TLR9 and contributing to disease pathogenesis^17–19^.

Imiquimod (IMQ), also known as R837, is an imidazoquinoline derivative with antiviral and immunomodulatory properties. IMQ-containing cream is clinically used to treat actinic keratoses, superficial basal cell carcinoma, and genital warts. However, repeated application of IMQ to mouse skin induces psoriasis-like dermatitis, providing a widely used experimental model of psoriasis^12^. Although IMQ-induced dermatitis reproduces several histological and inflammatory features of psoriasis and remains a widely used experimental model, the immune pathways driving IMQ-induced inflammation differ in part from those operating in human psoriasis^12,20^. Although IL-1β and inflammasome signaling can contribute to inflammatory responses in experimental models, their therapeutic relevance in plaque psoriasis remains less well established than that of the IL-23/IL-17 axis^21,22^. IMQ was originally characterized as a Toll-like receptor 7 (TLR7) agonist that activates MyD88-dependent NF-κB pathway, leading to proinflammatory cytokine production^23^. Beyond its canonical TLR7 signaling, IMQ also exerts TLR-independent effects^24,25^. It directly binds to mitochondrial complex I, promoting mitochondrial reactive oxygen species (ROS) production in DCs. These mitochondrial perturbations drive NLRP3 inflammasome activation, which mediates IL- 1β maturation and pyroptotic inflammation. Nevertheless, IMQ-induced skin inflammation remains only partially attenuated in TLR7- or NLRP3-deficient mice, suggesting that additional pathways contribute to IMQ-driven inflammatory responses ^26,27^. Accordingly, mechanistic findings obtained using IMQ should be interpreted in the context of this experimental model and subsequently validated for their relevance to human disease^20,26,28^.

The endoplasmic reticulum (ER) is essential for maintaining protein quality control, lipid metabolism, and intracellular calcium homeostasis^23^. In response to environmental insults such as oxidative stress, infection, or UV irradiation, ER function can be perturbed, triggering the unfolded protein response (UPR) through IRE1α, PERK, and ATF6 signaling. While primarily cytoprotective, sustained or dysregulated UPR signaling contributes to inflammation and has been implicated in chronic inflammatory diseases^29,30^. The ER also forms physical contact sites with mitochondria, referred to as mitochondria-associated ER membranes (MAMs), which play a critical role in calcium exchange and mitochondrial integrity^31^. Excessive or dysregulated ER-mitochondria tethering has been associated with mitochondrial calcium overload, ROS generation, and organelle dysfunction^31^. Previous studies have suggested a role for ER stress in concert with TLR in worsening psoriasis^32^. However, the upstream events linking IMQ to organelle stress responses remained unclear.

Gelsolin is a multifunctional, actin-binding protein that exists in two major isoforms: cytoplasmic (cGelsolin) and secreted plasma (pGelsolin)^33,34^. It regulates actin filament assembly and disassembly in a calcium-dependent manner and plays critical roles in cell morphology, migration, and signal transduction. Recent studies have highlighted its importance in immune regulation. Gelsolin has been implicated in the regulation of antigen cross-presentation by dendritic cells through its interaction with actin filaments^35^. Furthermore, it has been shown to suppress NLRP3 inflammasome activation by stabilizing cytosolic ion homeostasis and reducing mitochondrial stress^36^. Gelsolin also binds intracellular calcium and may contribute to cellular adaptation to calcium dysregulation during stress responses^33,34^. Recent transcriptomic and proteomic studies have reported altered Gelsolin expression in patients with psoriasis, suggesting a potential association between Gelsolin dysregulation and disease pathogenesis^37^. Despite emerging evidence linking altered Gelsolin expression to psoriasis, its functional role and its interaction with ER stress pathways remain poorly characterized.

Here, we investigated how IMQ induces ER stress and inflammasome-associated inflammatory responses in a psoriasis-like dermatitis model. Moreover, we identify Gelsolin as a regulator of UPR signaling and mitochondrial homeostasis during IMQ-induced inflammatory responses. Loss of Gelsolin exacerbates mitochondrial dysfunction, and its reduced expression correlates with psoriasis severity, highlighting both Gelsolin and UPRs as potential therapeutic targets.

## Results

### IMQ Induces Cytosolic and Mitochondrial Ca²^+^ flux, leading to Mitochondrial Dysfunction and Inflammasome Activation in DCs

To elucidate how IMQ activates innate immune signaling, we first examined intracellular Ca^2+^ dynamics in LPS-primed bone marrow-derived dendritic cells (BMDCs). Live-cell imaging with Fluo-8 revealed that IMQ, but not the related TLR7 agonist resiquimod (RSQ), induced a rapid increase in cytosolic Ca^2+^ concentration (Figure S1). This response was retained in MyD88-deficient BMDCs (Figure 1A), indicating a MyD88-independent mechanism. Flow cytometric analysis using the mitochondrial Ca²^+^ indicator Rhod-2 showed that IMQ also promoted Ca²^+^ transfer toward mitochondria (Figure 1B), suggesting that cytosolic Ca^2+^ elevation is accompanied by enhanced ER–mitochondria communication. IMQ stimulation led to an increase in intracellular reactive oxygen species (ROS) levels, whereas treatment with 2- APB, an inhibitor for IP_3_ receptor, significantly reduced ROS production (Figure 1C). Consistently, 2-APB also suppressed IL-1β release, cell death, and Caspase-1 cleavage (Figure 1D, 1E, and 1F), indicating that intracellular Ca^2+^ mobilization contributes to inflammasome activation and pyroptosis. In addition, IMQ-induced IL-1β production and cell death induction was abrogated by NLPR3 deficiency (Figure S2), demonstrating IMQ promotes NLRP3 inflammasome activation. To assess mitochondrial responses, we monitored mitochondrial membrane potential and mitochondrial ROS generation. IMQ, but not RSQ, caused depolarization and increased mitochondrial ROS levels (Figure 1G, and 1H). Treatment with the antioxidants glutathione ethyl ester (GSHEE), NAC, or PDTC significantly reduced IMQ-induced IL-1β release and cell death (Figure 1I, and 1J), and the mitochondrial-targeted antioxidant MitoQ showed a similar inhibitory effect (Figure 1K). Together, these results demonstrate that IMQ elevates intracellular and mitochondrial Ca²^+^ and ROS levels, both of which are closely associated with NLRP3 inflammasome activation in dendritic cells.

**Figure 1.**
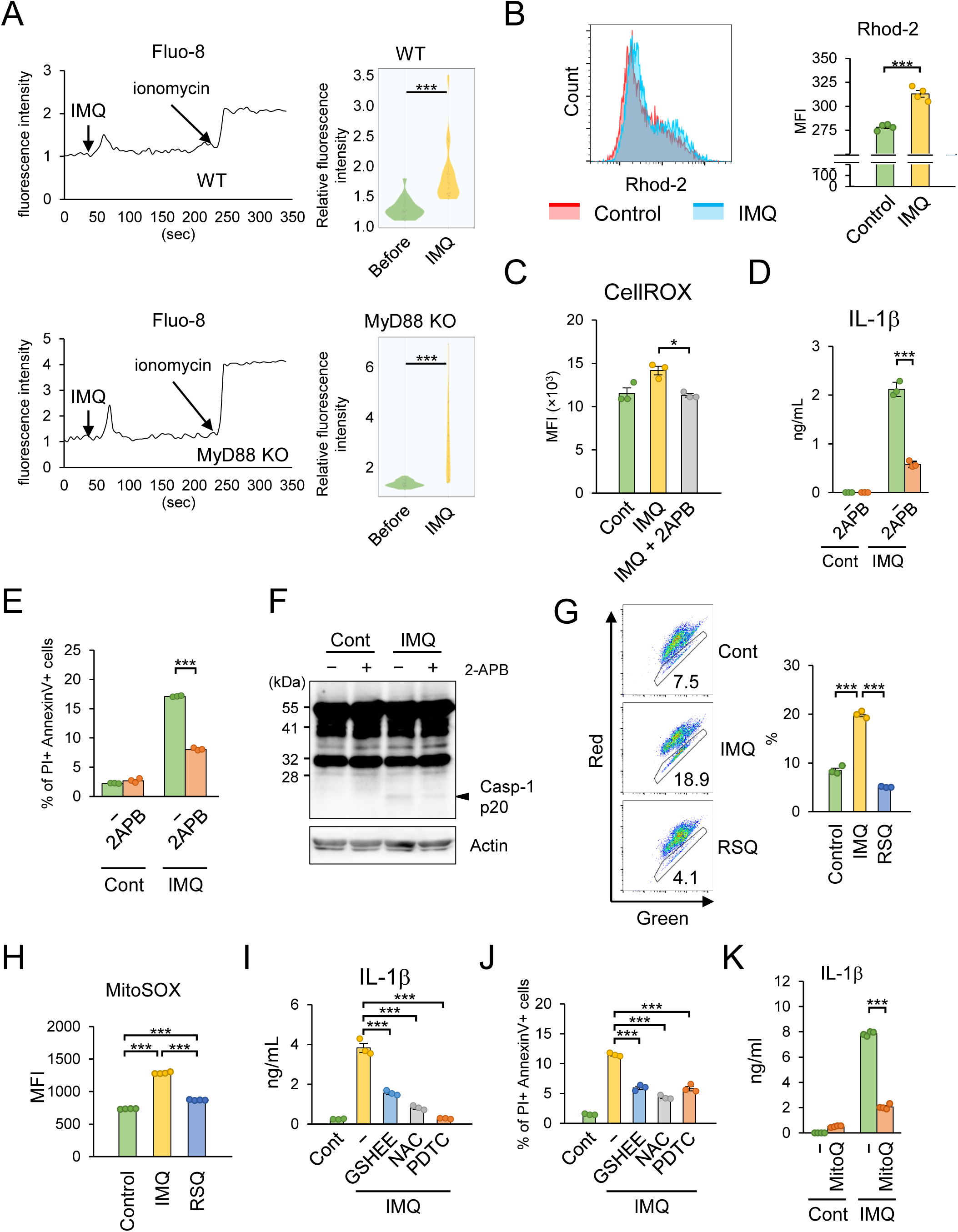
IMQ induces cytosolic and mitochondrial Ca^2+^ flux, leading to mitochondrial dysfunction and inflammasome activation in DCs. **a,** WT and MyD88-deficient BMDCs were pre-stained with Fluo-8, followed by IMQ stimulation. Ca^2+^ influx was monitored by fluorescent microscopy. Left panel shows representative Fluo-8 fluorescent intensity images. Right panel shows violin plots (*n* = 20). **b,** BMDCs were pre-stained with Rhod-2-AM reagent, followed by LPS stimulation. Mitochondrial Ca^2+^ was analyzed by flow cytometry (*n* = 4). Left panel shows a representative flow cytometry image. **c,** BMDCs were pretreated with 2-APB, followed by IMQ stimulation. Intracellular ROS was analyzed by flow cytometry (*n* = 3). **d, e, f,** BMDCs were pretreated with LPS, followed by 2-APB treatment, and stimulated with IMQ. IL-1β concentrations in cell culture supernatant were measured by ELISA (d) (*n* = 3). Cells were stained with PI and Annexin V, and cell death was analyzed by flow cytometry (e) (*n* = 3). Cell lysates were analyzed by immunoblotting for cleaved caspase- 1 (p20) (f). Data are representative immunoblot analysis images of two independent experiments. **g,** BMDCs were stimulated with IMQ or RSQ, stained with JC-1 reagent, and mitochondrial damage was analyzed by flow cytometry (*n* = 3). Left panel shows representative flow cytometry images. **h,** BMDCs were pre-stained with MitoSOX red reagent, stimulated with IMQ or RSQ, and mitochondrial ROS was analyzed by flow cytometry (*n* = 4). Left panel shows representative flow cytometry images. **i,** BMDCs were pretreated with LPS, followed by treatment with the indicated anti-oxidants, stimulated with IMQ, and IL-1β concentrations in cell culture supernatants were measured by ELISA (*n* = 3). **j,** BMDCs were pretreated with LPS, followed by treatment with the indicated anti-oxidants, stimulated with IMQ, stained with PI and Annexin V, and cell death was analyzed by flow cytometry (*n* = 3). **k,** BMDCs were pretreated with mitoquinone (MitoQ), followed by IMQ stimulation, and IL-1β concentrations in cell culture supernatants were measured by ELISA (*n* = 3). Data are presented as the mean ± s.e.m. Each circle indicates an independent biological sample. *P-*values were calculated by one-way ANOVA with Tukey’s test (c-e and g-k) or Student’s *t*-test (a and b). (*: *p* < 0.05, ***: *p* < 0.001).

### IMQ Promotes UPR and Facilitates IL-23 Expression in DCs

Given that IMQ increased cytosolic and mitochondrial Ca^2+^ and promoted ER– mitochondria communication, we next examined whether ER homeostasis is affected by IMQ stimulation. A proximity ligation assay revealed that IMQ enhanced the formation of MAMs, indicating strengthened ER–mitochondria contacts (Figure 2A). To elucidate the role of IMQ in modulating ER bioprocesses, we investigated its impact on unfolded protein response (UPR), a key cellular mechanism activated under ER stress, by quantifying the expression of UPR-related markers. We first examined the spliced form of *Xbp1* mRNA (*Xbp1s* mRNA), a critical indicator of ER stress generated through IRE1α. IMQ stimulation significantly upregulated *Xbp1s* mRNA expression in both WT and MyD88-deficient BMDCs, demonstrating robust induction of ER stress through a MyD88-independent pathway (Figure 2B). This finding suggests that IMQ triggers ER stress independently of the canonical TLR signaling. By contrast, RSQ stimulation did not induce *Xbp1s* mRNA expression in BMDCs, indicating that the ER stress response is specific to IMQ and not a general feature of TLR7/8 agonists (Figure S3A). To confirm the involvement of IRE1α in IMQ-induced *Xbp1s* expression, we pretreated BMDCs with 4μ8c, a selective inhibitor of IRE1α RNase activity. This pretreatment significantly suppressed IMQ-induced *Xbp1s* mRNA expression, underscoring the critical role of IRE1α in mediating this ER stress response (Figure S3B).

**Figure 2.**
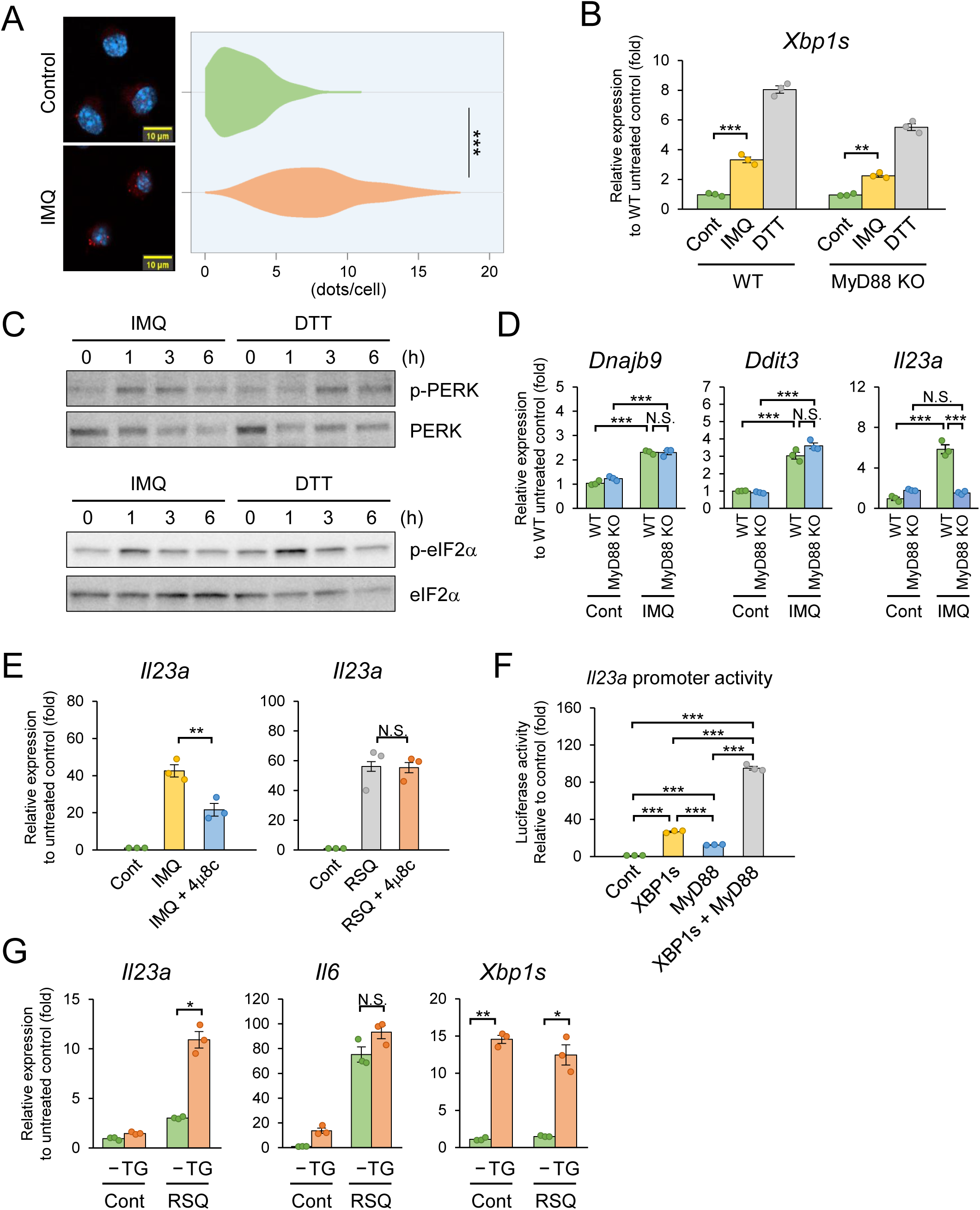
IMQ promotes UPR and facilitates *Il23a* expression in DCs. **a,** BMDCs were pretreated with LPS, followed by IMQ stimulation, and VDAC1-IP_3_R1 interactions were analyzed by PLA. Left panel shows representative PLA images (*n* = 200). **b,** WT and MyD88-deficient BMDCs were stimulated with IMQ or DTT as a positive control, and the expression of short form *Xbp1* (*Xbp1s*) mRNA was measured by qPCR (*n* = 3). **c,** BMDCs were stimulated with IMQ or DTT as a positive control, and cell lysates were analyzed by immunoblotting for the phosphorylated form of PERK and eIF2α. Data are representative images of two independent experiments. **d,** WT and MyD88-deficient BMDCs were stimulated with IMQ, and the expressions of *Dnajb9, Ddit3,* and *Il23a* mRNA were measured by qPCR (*n* = 3). **e,** BMDCs were pretreated with IRE1α ribonuclease inhibitor 4μ8c, followed by IMQ or RSQ stimulation, and the expression of *Il23a* mRNA was measured by qPCR (*n* = 3). **f,** HEK293T cells were transfected with an *Il23a* promoter reporter plasmid together with the indicated expression plasmids, and luciferase activity was measured (*n* = 3). **g,** BMDCs were stimulated with RSQ and thapsigargin, and the expressions of *Il23a*, *Il6,* and *Xbp1s* mRNA were measured by qPCR (*n* = 3). Data are presented as the mean ± s.e.m. Each circle indicates an independent biological sample. *P-*values were calculated by one-way ANOVA with Tukey’s test (b and d-g) or Student’s *t*-test (a). (N.S.: not significant, *: *p* < 0.05, **: *p* < 0.01, ***: *p* < 0.001).

In addition to the IRE1α-Xbp1 axis, we explored other branches of the UPR to gain a comprehensive understanding of effects of IMQ on ER stress signaling. IMQ stimulation markedly increased the phosphorylation of PERK and its downstream target, eIF2α, in BMDCs, but not RSQ (Figure 2C, and S3C). Furthermore, IMQ treatment upregulated the expression of UPR-associated genes, including *Dnajb9* (also known as *Erdj4*) and *Ddit3* (also known as *Chop*), which are hallmarks of ER stress responses, in a Myd88-independent manner (Figure 2D). Collectively, these results demonstrate that IMQ potently induces multiple UPR pathways through ER stress, including IRE1α and PERK, independently of MyD88 signaling.

Previous studies have reported that ROS induces ER stress^38,39^. To investigate whether mitochondria are required for ER stress induction upon IMQ stimulation, we examined the effects of mitochondrial depletion. Treatment with the mitochondrial uncoupler carbonyl cyanide m-chlorophenyl hydrazone (CCCP) combined with ectopic expression of the ubiquitin E3 ligase Parkin induces robust mitophagy, leading to effective mitochondrial clearance from cells. We generated Parkin-expressing RAW264.7 cells and cultured them with CCCP to induce mitochondrial depletion (Figure S4A and S4B). In both WT and mitochondria-depleted cells, IMQ stimulation increased the expression of ER stress-related genes (Figure S4C). These findings suggest that IMQ activates ER stress through a mechanism that is distinct from its effects on mitochondria. Since IL-23 is a critical inflammatory cytokine in the pathogenesis of psoriasis^3,40,41^, we next evaluated the impact of IMQ treatment on *Il23a* expression in BMDCs. IMQ treatment markedly induced *Il23a* expression (Figure 2D), whereas this induction was completely abolished in MyD88-deficient cells. Interestingly, pretreatment with 4μ8c significantly suppressed IMQ-induced *Il23a* transcription but not RSQ- induced *Il23a* transcription (Figure 2E), suggesting that UPR enhances IMQ-mediated *Il23a* expression. A luciferase assay using an *Il23a* promoter construct (−2,030 to +20 bp) revealed that *Il23a* promoter activity was increased by XBP1s or MyD88 overexpression and synergistically enhanced by their co-expression (Figure 2F, and S5). Additionally, RSQ-induced *Il23a* expression, which was induced in an IRE1α-independent manner, was enhanced by thapsigargin (Tg), an ER stress inducer, whereas *Il6* expression was unaffected (Figure 2G). These results suggest that IMQ activates TLR and ER stress pathways via distinct mechanisms, which converge to cooperatively enhance *Il23a* expression.

### IMQ Drives Psoriasis-Associated Gene Expression in Keratinocytes via the IRE1α- XBP1s Axis of the UPR

In psoriatic lesions, keratinocytes exhibit hyperproliferation and aberrant differentiation, contributing to the inflammatory milieu of psoriasis^1,2^. To investigate whether IMQ induces UPRs in keratinocytes, we stimulated primary mouse keratinocytes with IMQ and assessed key UPR markers. RT-qPCR analysis revealed that IMQ significantly upregulated mRNA expression of *Xbp1s*, *Hspa5*, and *Ddit3*, indicating robust activation of the UPRs (Figure 3A). Concurrently, IMQ stimulation increased cytoplasmic Ca²^+^ influx, as measured by Fluo-8 staining, and enhanced phosphorylation of PERK and eIF2α, further confirming UPR activation (Figure 3B, and 3C).

**Figure 3.**
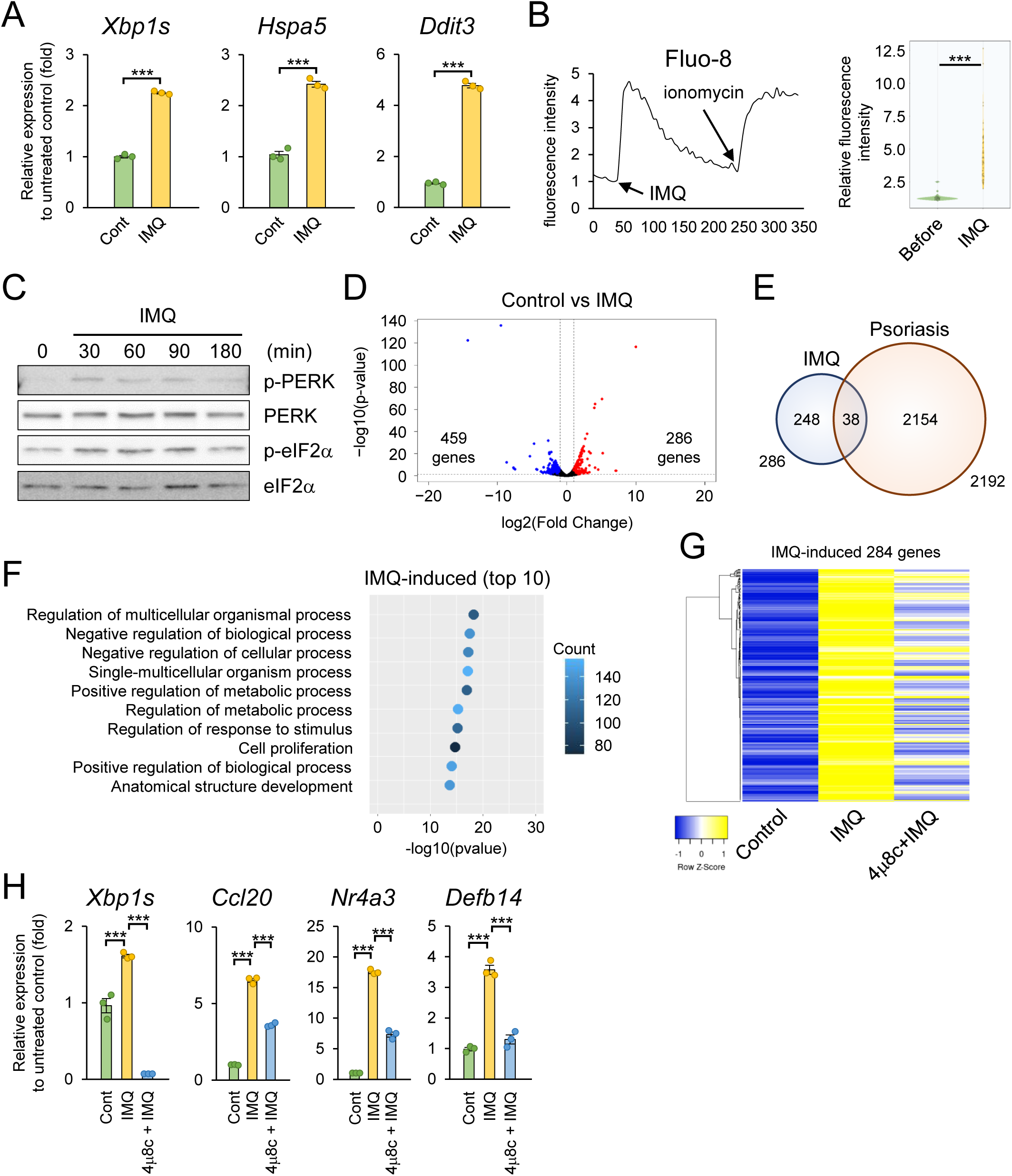
IMQ drives psoriasis-associated gene expression in keratinocytes via the IRE1α-XBP1s axis of the UPR. **a,** Mouse keratinocytes were stimulated with IMQ, and the expressions of *Xbp1s*, *Hspa5,* and *Ddit3* mRNA were measured by qPCR (*n* = 3). **b,** Mouse keratinocytes were pre-stained with Fluo-8, followed by IMQ stimulation. Ca^2+^ influx was monitored by fluorescent microscopy (*n* = 40). **c,** Mouse keratinocytes were stimulated with IMQ, and cell lysates were analyzed by immunoblotting for the phosphorylated form of eIF2α and PERK. Data are representative images of two independent experiments. **d,** Mouse keratinocytes were stimulated with IMQ, and the gene expression profile was analyzed by RNA-seq. Data are shown as a volcano plot. Red and blue plots represent significantly upregulated and downregulated genes (*p* < 0.05, |log2FoldChange| > 1), respectively. **e,** Upregulated genes in (d) were compared with those in skin from human psoriasis patients. Data are shown as a Venn diagram. **f,** Upregulated genes in (d) were subjected to GO analysis (BP2). Data shows the top 10 (*p-*value) enriched biological processes. **g,** Data shows a heat map of genes upregulated in (d). **h,** Mouse keratinocytes were pretreated with 4μ8c, followed by IMQ stimulation, and the expressions of *Xbp1s*, *Ccl20*, *Nr4a3,* and *Defb14* mRNA were measured by qPCR (*n* = 3). Data are presented as the mean ± s.e.m. Each circle indicates an independent biological sample. *P-*values were calculated by one-way ANOVA with Tukey’s test (h) or Student’s *t*-test (a and b). (***: *p* < 0.001).

To explore the broader impact of IMQ-induced UPRs on keratinocyte gene expression, we performed RNA-sequencing (RNA-seq) analysis on IMQ-stimulated primary mouse keratinocytes, with or without pretreatment with 4μ8c. Differential expression analysis identified 286 upregulated and 459 downregulated genes in IMQ-treated keratinocytes compared to controls (Figure 3D, and Tables S1, and S2). Notably, 38 (13%) of the upregulated genes overlapped with those previously reported in human psoriatic lesions, suggesting that IMQ recapitulates key molecular features of psoriasis in keratinocytes (Figure 3E, and Tables S3, and S4). Gene ontology (GO) analysis of the upregulated genes revealed significant enrichment of pathways related to cellular metabolism and proliferation, consistent with the hyperproliferative phenotype of psoriatic keratinocytes (Figure 3F, and Table S5). Pretreatment with 4μ8c significantly reversed IMQ-induced gene expression changes, indicating that the IRE1α-XBP1s axis is critical for these transcriptional responses (Figure 3G, and Table S6).

To validate these findings, we performed RT-qPCR on selected psoriasis-associated genes, including *Ccl20*, *Nr4a3*, and *Defb14* (the mouse ortholog of human β- defensin 3, hBD3), which were selected because our transcriptomic analysis identified them as representative UPR-responsive genes strongly induced by IMQ stimulation. IMQ significantly induced the expression of these genes, and 4μ8c pretreatment effectively reversed this induction, confirming the dependency on IRE1α-mediated UPR signaling (Figure 3H). To further assess the relevance of these findings to psoriasis-associated inflammatory signaling, primary mouse keratinocytes were stimulated with IL-17A and TNF-α. As expected, IL-17A and TNF-α markedly induced the expression of canonical psoriasis-associated genes, including S100a8, S100a9, and Defb14, the latter of which was also induced by IMQ stimulation (Figure S6). These findings demonstrate that IMQ activates the UPR in keratinocytes, particularly through the IRE1α-XBP1s axis, to drive the expression of psoriasis-associated genes. Furthermore, the induction of canonical psoriasis-associated genes by IL-17A and TNF-α supports the relevance of these findings to psoriasis-associated inflammatory signaling.

### UPRs Exacerbate IMQ-Induced Dermatitis, and Ca^2+^ Flux Inhibition Ameliorates It

To investigate the in vivo role of the UPR in IMQ-induced psoriasis-like dermatitis, we evaluated the expression of ER stress-related genes in a mouse model. We quantified mRNA levels of key UPR markers in ear tissue following topical IMQ application. Our results revealed significant upregulation of *Xbp1s*, *Hspa5* (also known as BiP or Grp78), and *Hsp90b1* (also known as Grp94) mRNA in IMQ-treated ears compared to untreated controls, indicating robust induction of ER stress in psoriatic lesions (Figure 4A).

**Figure 4.**
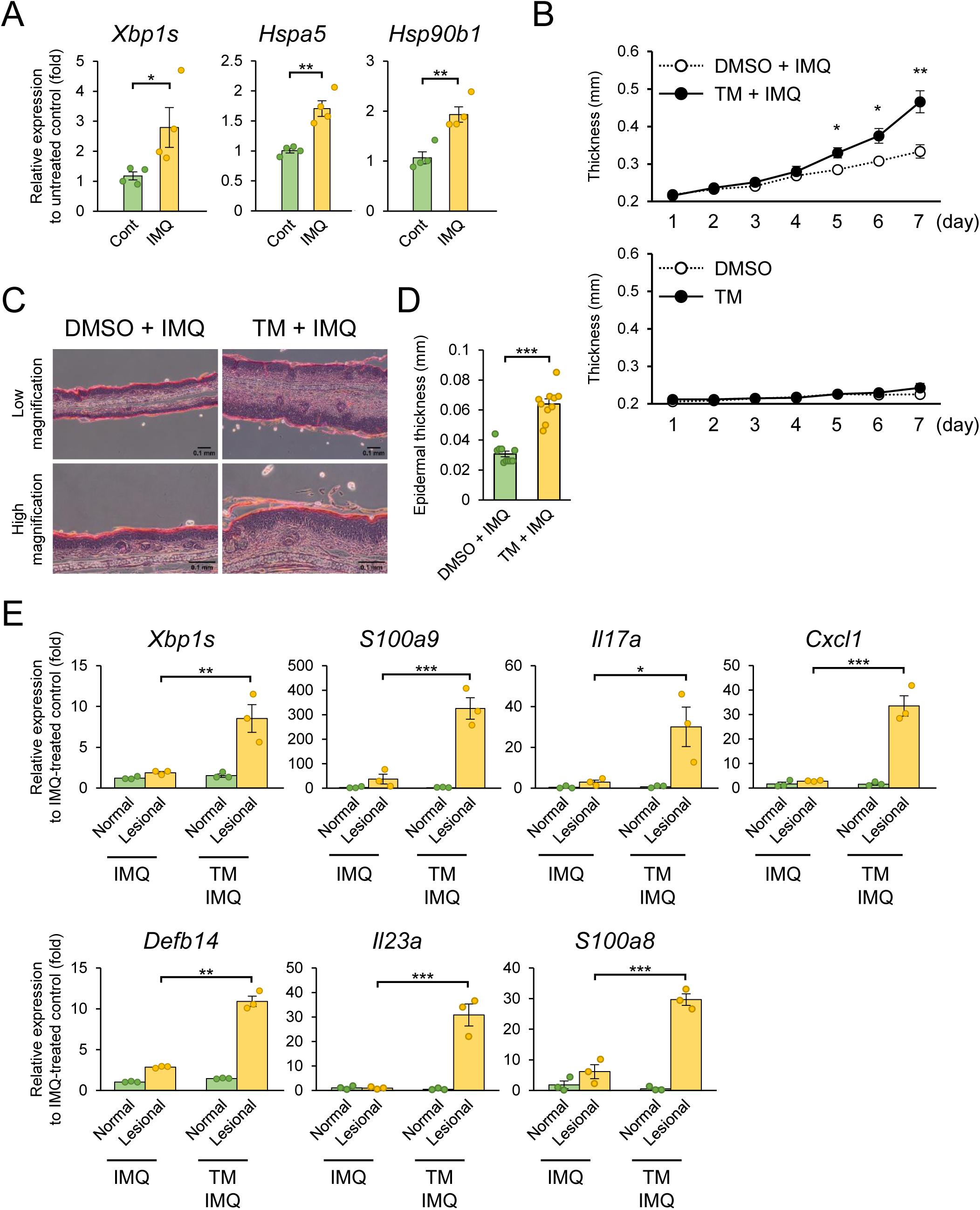
UPRs exacerbate IMQ-induced dermatitis, and Ca^2+^ flux inhibition ameliorates it. **a,** IMQ cream was applied to the ear lobes of mice. Total RNA was extracted from the ear lobes and the expressions of *Xbp1s*, *Hspa5,* and *Hsp90b1* mRNA were measured by qPCR (*n* = 4). **b, c, d, e,** IMQ cream and tunicamycin, an ER stress inducer, were applied daily to the ear lobes of mice. Ear thickness was measured daily (*n* = 3) (c). Mouse ears were sampled and subjected to histological analysis (H&E staining) (d). The epidermal thickness in (d) was measured (*n* = 10) (e). Total RNA was harvested from the ears, and the expression of *Xbp1s*, *S100a9*, *Il17a*, *Cxcl1,* and *Defb14* mRNA was measured by qPCR (*n* = 3) (f). Data are presented as the mean ± s.e.m. Each circle indicates an independent biological sample. *P-*values were calculated by one-way ANOVA with Tukey’s test (f) or Student’s *t*-test (a-c and e). (N.S.: not significant, *: *p* < 0.05, **: *p* < 0.01, ***: *p* < 0.001).

To further elucidate the functional role of UPR in IMQ-induced dermatitis, we pretreated mice with tunicamycin, a pharmacological inducer of ER stress, prior to IMQ application. Tunicamycin pretreatment markedly exacerbated IMQ-induced ear thickening and epidermal hyperplasia, as assessed by histological analysis (Figure 4B, 4C, and 4D). Additionally, tunicamycin significantly enhanced the expression of psoriasis-associated genes, including the canonical psoriasis signature genes *S100a8* and *Il23a*, as well as *Defb14* (encoding mouse BD14 [mBD14]), in IMQ-treated ears, indicating that heightened ER stress amplifies the inflammatory and psoriatic phenotype (Figure 4E, and S7). To examine the role of calcium signaling in this context, we treated mice with carbachol, an inositol 1,4,5-trisphosphate receptor (IP_3_R) activator that promotes Ca^2+^ release from the ER. Carbachol treatment significantly worsened IMQ-induced dermatitis, as evidenced by increased ear thickening, epidermal hyperplasia and enhanced expression of psoriasis-related inflammatory genes, further supporting the role of Ca^2+^-mediated ER stress in disease exacerbation (Figure S8).

Conversely, to assess whether inhibiting Ca^2+^ flux could reduce IMQ-induced pathology, we pretreated mice with 2-APB. Strikingly, 2-APB pretreatment significantly reduced IMQ-induced ear thickening and epidermal hyperplasia, as determined by histological analyses (Figure 5A, 5B, and 5C). Moreover, 2-APB treatment markedly decreased the expression of inflammatory and psoriasis-related genes in IMQ-treated ears (Figure 5D). Collectively, these findings demonstrate that Ca^2+^-triggered ER stress, coupled with mitochondrial damage, plays a pivotal role in driving IMQ-induced psoriasis-like dermatitis. Furthermore, inhibition of Ca^2+^ flux via IP_3_R blockade ameliorates disease severity, highlighting a potential therapeutic strategy for targeting ER stress in psoriasis.

**Figure 5.**
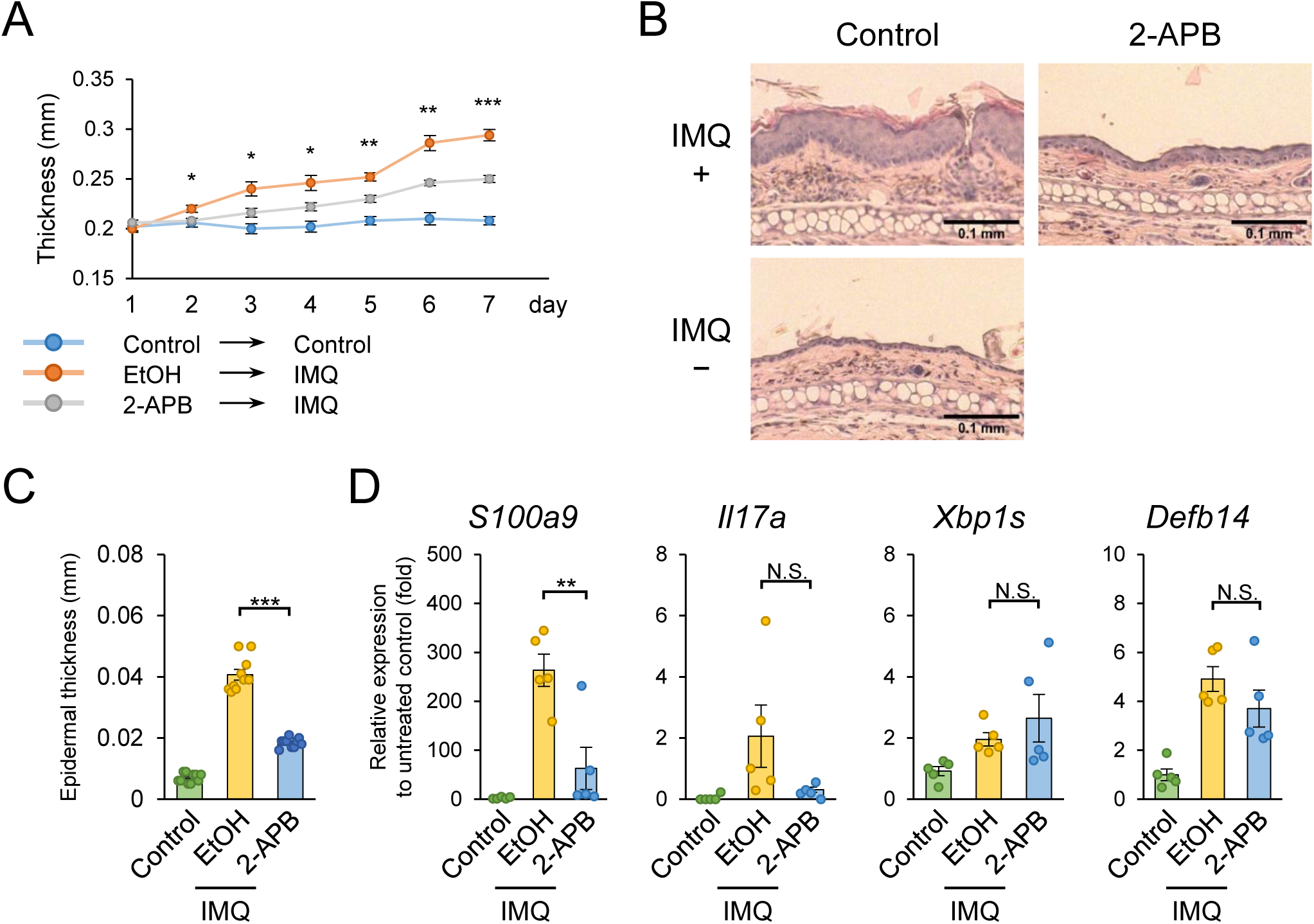
Ca^2+^ flux inhibition ameliorates dermatitis. **a, b, c, d,** IMQ cream and 2-APB were applied daily to the ear lobes of mice. Ear thickness was measured daily (*n* = 5) (a). Mouse ears were sampled and subjected to histological analysis (H&E staining). Data are shown as representative histological analysis images (b). The epidermal thickness in (b) was measured (*n* = 10) (c). Total RNA was harvested from the ear and the expression of *S100a9*, *Il17a*, *Xbp1s,* and *Defb14* mRNA was measured by qPCR (*n* = 5) (d). Data are presented as the mean ± s.e.m. Each circle indicates an independent biological sample. *P-*values were calculated using one-way ANOVA with Tukey’s test. (N.S.: not significant, *: *p* < 0.05, **: *p* < 0.01, ***: *p* < 0.001).

### Mitochondrial DNA and mBD14 Complex Activate pDC via TLR9

Mitochondrial damage promotes the release of mtDNA into the cytoplasm, where it eventually activates the NLRP3 inflammasome^42,43^. We found that IMQ stimulation significantly increased mtDNA levels in both mitochondrial and cytoplasmic fractions, while cytosolic nuclear DNA (nDNA) remained unchanged (Figure 6A). Furthermore, 2-APB treatment significantly suppressed increment of cytoplasmic mtDNA induced by IMQ (Figure 6B). IMQ also increased the amount of oxidized DNA in cytosolic fraction, as measured by dot blot analysis (Figure 6C), and enhanced the presence of mitochondria in the extracellular milieu (Figure 6D). Furthermore, extracellular mtDNA levels were elevated in an NLRP3-dependent manner (Figure 6E). These results suggest that IMQ increases cytoplasmic mtDNA levels, which contribute to NLRP3 inflammasome-dependent pyroptosis and its subsequent release into the extracellular milieu.

**Figure 6.**
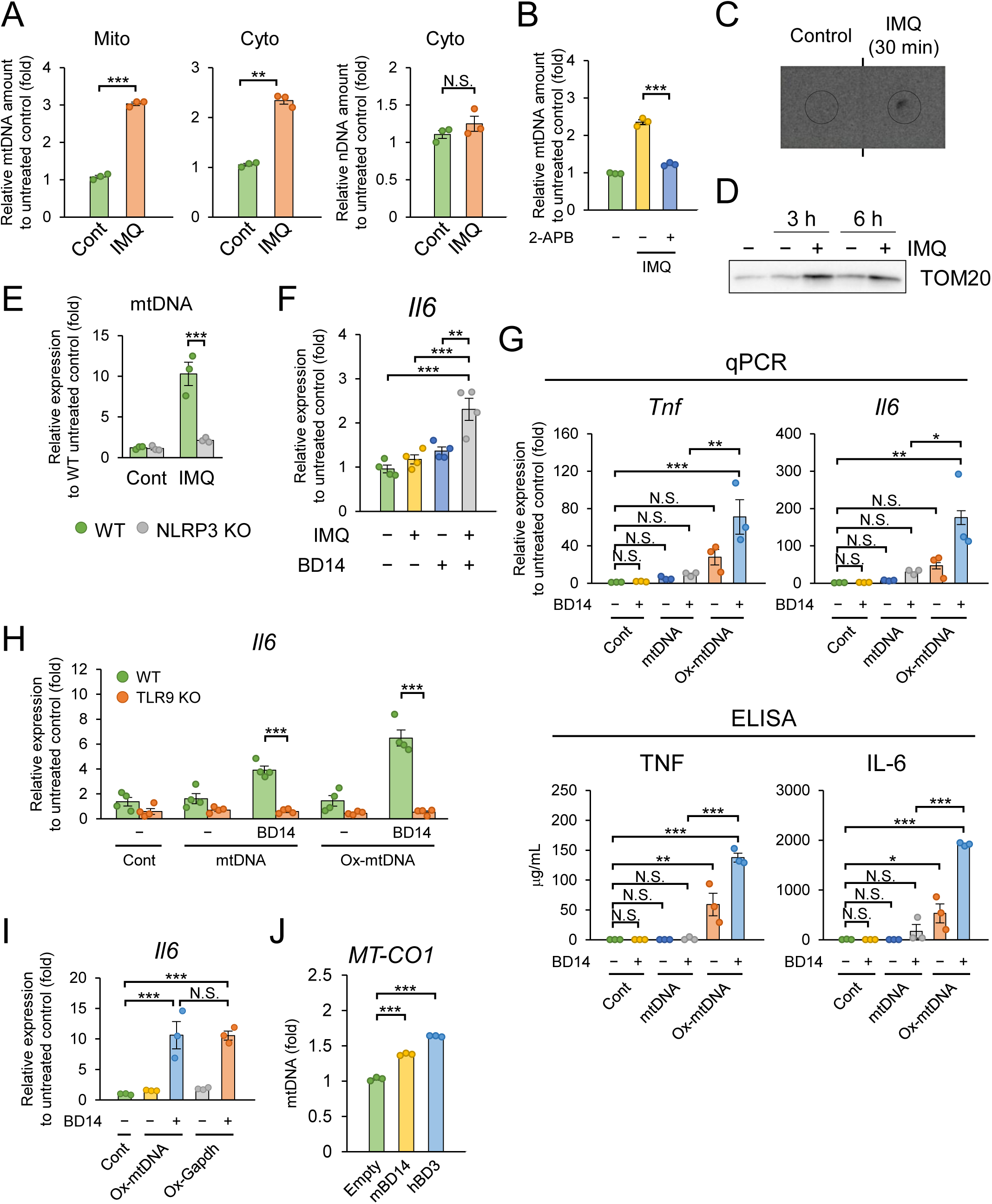
Mitochondrial DNA and mBD14 complex activate pDC via TLR9. **a,** BMDCs were pretreated with LPS, followed by IMQ stimulation. DNA was harvested from cytosolic and mitochondrial fractions, and the amount of mitochondrial DNA and nuclear DNA (nDNA) were measured by qPCR (*n* = 3). **b,** BMDCs were pretreated with LPS and 2-APB, followed by IMQ stimulation. DNA was harvested from cytosolic fraction, and the amount of mitochondrial DNA was measured by qPCR (*n* = 3). **c,** BMDCs were pretreated with LPS, followed by IMQ stimulation. DNA was harvested from cytosolic fraction, and oxidized DNA was detected by dot blot analysis. Data are representative images of two independent experiments. **d,** BMDCs were stimulated with IMQ, and mitochondria-enriched fractions were collected from the cell culture supernatants and lysed. Western blot analysis was performed using an anti-TOM20 antibody to visualize mitochondrial content in the extracellular fraction. Data are representative images of two independent experiments. **e,** WT and NLRP3-deficient BMDCs were pretreated with LPS, followed by IMQ stimulation. DNA in cell culture supernatants was harvested and the amount of mitochondrial DNA was measured by qPCR (*n* = 3). **f,** Bone marrow-derived pDCs were stimulated with DNA isolated from the cytosolic fraction of BMDCs stimulated with IMQ, and mBD14. Total RNA was extracted from the pDCs and the expression of *Il6* and *Tnf* mRNA was measured by qPCR (*n* = 4). **g,** BMDCs were stimulated with synthesized mtDNA or oxidized mtDNA with or without recombinant mBD14. The expression of *Tnf* and *Il6* mRNA was measured by qPCR (*n* = 3) (upper panel). The concentration of TNF-α and IL-6 in cell culture supernatants was measured by ELISA (*n* = 3) (bottom panel). **h,** WT and TLR9-deficient bone marrow-derived pDCs were stimulated with mtDNA or oxidized mtDNA with or without mBD14. The expression of *Il6* mRNA was measured by qPCR (*n* = 3). **i**, BMDCs were stimulated with oxidized double stranded-DNA synthesized from mitochondrial or genomic DNA. The expression of *Il6* mRNA was measured by qPCR (*n* = 3). **j,** HEK293T cells were transfected with FLAG-mBD14 or FLAG-hBD3 expression plasmids. Cell lysates were subjected to DNA immunoprecipitation with anti-FLAG antibody and immunoprecipitants were analyzed by qPCR (*n* = 3). Data are presented as the mean ± s.e.m. Each circle indicates an independent biological sample. *P-*values were calculated by one-way ANOVA with Tukey’s test (e-j) or Student’s *t*-test (a and b). (N.S.: not significant, *: *p* < 0.05, **: *p* < 0.01, ***: *p* < 0.001).

AMPs such as LL37 and β-defensins contribute to psoriasis pathogenesis by forming complexes with self-derived nucleic acids and promoting pDC activation through nucleic acid-sensing pathways including TLR9^17,18,40^. We hypothesized that DNA released from IMQ-stimulated DCs forms a complex with mBD14. Indeed, pDCs incubated with DNA from IMQ-stimulated BMDCs and recombinant mBD14 exhibited significantly higher *Il6* mRNA expression than those treated with DNA or mBD14 alone (Figure 6F). In contrast, DNA from unstimulated BMDCs failed to induce *Il6* expression, even in the presence of mBD14 (Figure 6F).

Since oxidized mtDNA activates NLRP3 in the cytoplasm^42,43^, we further hypothesized that it also enters the extracellular space and, together with mBD14 secreted by keratinocytes, activates pDCs. To test this, we treated pDCs with synthetic mtDNA (syn-mtDNA) with or without recombinant mBD14 and measured cytokine expression. Syn-mtDNA alone, even with mBD14, did not induce TNF or IL-6 expression (Figure 6G). However, co-treatment with the oxidized form (syn-ox-mtDNA) and mBD14 significantly upregulated TNF and IL-6 at both mRNA and protein levels (Figure 6G). This induction was abolished in TLR9- and MyD88-deficient pDCs, confirming the essential role of TLR9 in this response (Figure 6H). To determine whether DNA sequence specificity affects pDC activation, we tested synthetic oxidized double-stranded DNA from genomic DNA (syn-ox-Gapdh). Syn-ox-Gapdh induced *Il6* mRNA expression at levels comparable to syn-ox-mtDNA, suggesting that DNA oxidation, rather than sequence specificity, drives pDC activation (Figure 6I). Lastly, to assess mBD14-DNA interactions under physiological conditions, we overexpressed FLAG-tagged mBD14 or hBD3 in HEK293T cells and performed DNA immunoprecipitation using an anti-FLAG antibody. mtDNA was efficiently immunoprecipitated with both mBD14 and hBD3 (Figure 6J).

### IMQ Interacts with Gelsolin and Gelsolin Deficiency Increases UPR

To uncover the molecular mechanisms by which IMQ induces ER stress and activates UPRs, we sought to identify proteins that interact with IMQ (Figure 7A). IMQ was conjugated to magnetic beads and incubated with whole cell lysates from BMDCs. We next performed two rounds of mass spectrometric analysis and compiled a list of proteins that were specifically detected in the IMQ-beads, as well as those that were more abundantly detected in the IMQ-beads compared to the control beads (Table S7, and S8). Among these, we focused on Gelsolin, which was consistently detected in both independent experiments (Table S9). To validate this interaction, we overexpressed FLAG-tagged Gelsolin in HEK293T cells and incubated cell lysates with IMQ- conjugated beads. Immunoprecipitation assays confirmed that FLAG-Gelsolin was co-precipitated with IMQ-conjugated beads, but not with control beads (Figure 7B). Surface plasmon resonance (SPR) analysis further revealed a binding affinity (K_D) of approximately 1.8 μM, supporting the specificity of this interaction (Figure S9). These findings establish Gelsolin as a novel molecular interactor of IMQ, potentially regulating its proinflammatory effects.

**Figure 7.**
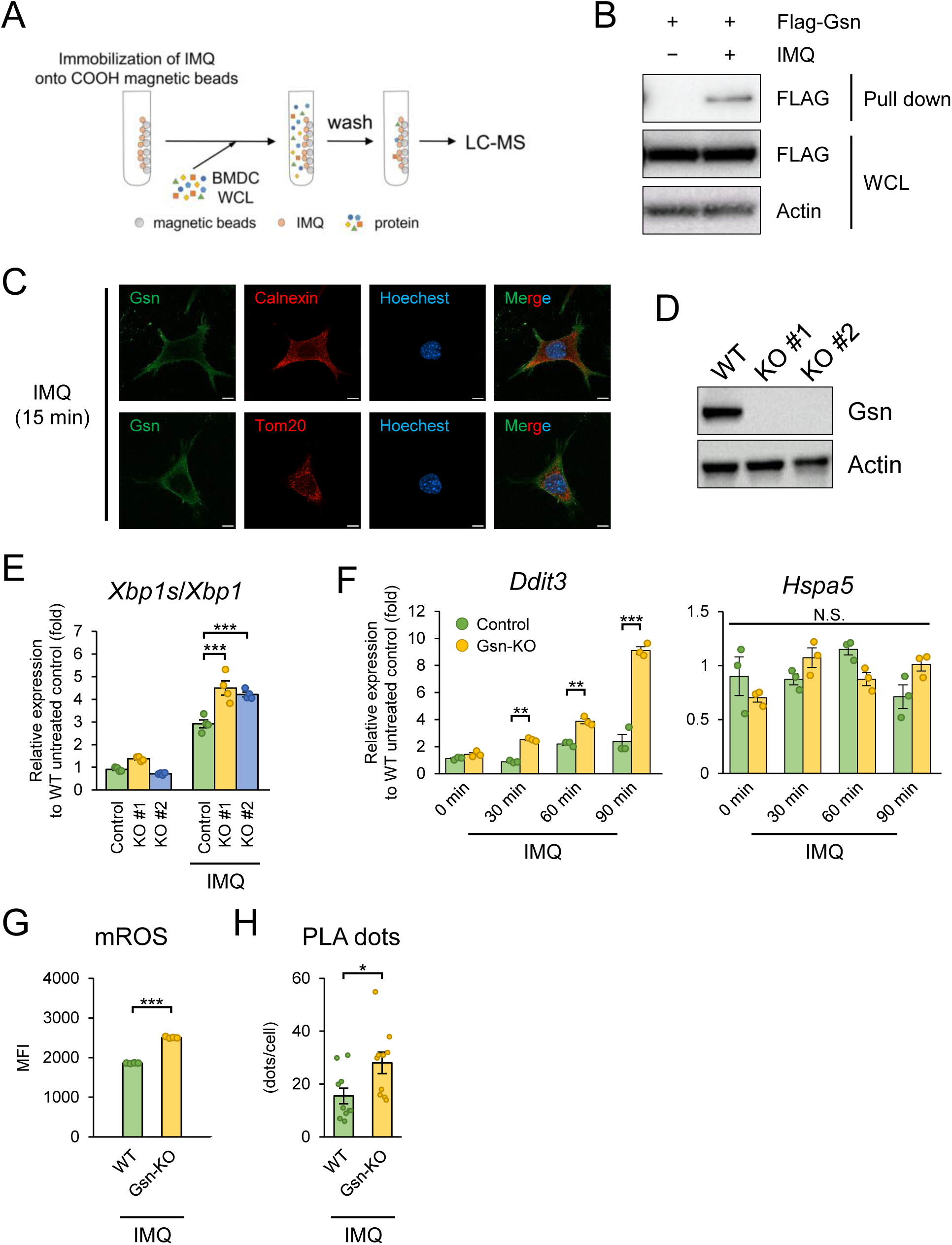
IMQ interacts with Gelsolin and Gelsolin deficiency increases UPRs. a, Schematic representation of the method to identify IMQ-binding proteins. **b,** IMQ-conjugated or control magnetic beads were incubated with whole cell lysate prepared from Gelsolin-overexpressed HEK293T cells. After pull-down, proteins were eluted from each bead, and subjected to western blot analysis for FLAG-Gelsolin. Data are representative images of two independent experiments. **c,** FLAG-Gelsolin was transiently expressed in MEFs and cells were stimulated with IMQ. Gelsolin was visualized in green, Calnexin (ER) and TOM20 (mitochondria) were stained in red, and nuclei were counterstained with Hoechst. Data are representative images of two independent experiments. Scale bars represent 10 μm. **d,** WT and Gelsolin (Gsn)-deficient MEFs were lysed, and cell lysates were analyzed by western blot analysis for Gelsolin expression. Data are representative images of two independent experiments. **e, f,** WT (Control) and Gsn-deficient MEFs were stimulated with IMQ, and *Xbp1s* (e), *Ddit3* (f) and *Hspa5* (f) expression was measured by qPCR analysis (e: *n* = 4, f: *n* = 3) **g,** WT and Gsn-deficient MEFs were pre-stained with MitoSOX red reagent, stimulated with IMQ, and mitochondrial ROS was analyzed by flow cytometry (*n* = 4). **h,** WT and Gsn-deficient MEFs were pretreated with LPS, followed by IMQ stimulation, and VDAC1-IP_3_R1 interactions were analyzed by PLA. (*n* = 10). Data are presented as the mean ± s.e.m. Each circle indicates an independent biological sample. *P-*values were calculated by one-way ANOVA with Tukey’s test (e and f) or Student’s *t*-test (g and h). (N.S.: not significant, *: *p* < 0.05, **: *p* < 0.01, ***: *p* < 0.001).

To investigate the subcellular localization of Gelsolin and its potential dynamics upon IMQ stimulation, we expressed FLAG-tagged Gelsolin in mouse embryonic fibroblast (MEF) and performed immunostaining before and after IMQ treatment. Under basal conditions, FLAG-Gelsolin did not colocalize with both ER and mitochondria (Figure S10). Following IMQ stimulation, however, we observed a portion of FLAG- Gelsolin colocalized with both ER and mitochondria (Figure 7C). These findings suggest that Gelsolin relocalizes to ER and mitochondria in response to IMQ and may function at ER to regulate stress responses.

To investigate the role of Gelsolin in IMQ-induced ER stress, we generated Gelsolin-deficient MEFs using the CRISPR/Cas9 system (Figure S11), achieving complete knockout as confirmed by western blot analysis (Figure 7D). Gelsolin-deficient MEFs and WT control cells were stimulated with IMQ, and the UPR was assessed. RT-qPCR revealed significantly elevated mRNA expression of *Xbp1s* and *Ddit3* in Gelsolin- deficient MEFs compared to WT cells following IMQ stimulation, indicating enhanced UPR activation (Figure 7E, and 7F). Given the interplay between ER stress and mitochondrial function, we examined mitochondrial parameters in IMQ-treated MEFs. MitoSOX staining revealed significant increase in mitochondrial ROS production in Gelsolin-deficient MEFs compared to WT cells (Figure 7G). Moreover, PLA assay revealed that IMQ stimulation increased ER-mitochondria contact sites in Gelsolin- deficient MEFs compared to WT cells (Figure 7H), indicating enhanced ER-mitochondria tethering.

To determine whether these effects were specific to IMQ or reflected a general consequence of TLR7 activation, Gelsolin-deficient MEFs were also stimulated with RSQ. In contrast to IMQ stimulation, RSQ did not consistently enhance Xbp1s expression, mitochondrial ROS production, or MAM formation in Gelsolin-deficient MEFs compared with WT cells (Figure S12). These findings indicate that the effects of Gelsolin deficiency are more pronounced during IMQ stimulation than during TLR7 activation induced by RSQ.

Collectively, these results suggest that Gelsolin suppresses IMQ-induced ER stress and mitochondrial dysfunction by modulating ER-mitochondria interactions, thereby exerting a protective role in psoriasis pathogenesis.

### Human and Mouse Psoriatic Lesions Exhibit Reduced Gelsolin and Enhanced ER Stress

To further explore the physiological link between Gelsolin, psoriasis pathogenesis, and the UPR, we re-analyzed RNA-seq datasets derived from non-lesional and lesional skin in a mouse model of IMQ-induced psoriasiform dermatitis, as well as from normal and lesional skin of human psoriasis patients. Re-analysis of the RNA-seq datasets revealed a set of transcriptional alterations characteristic of psoriatic lesions. Both in mice and humans, the expression of psoriasis-associated genes, including *Il23a* and *S100a9*, was markedly elevated in lesional skin (Figure 8A and 8B). In human lesions, *Defb103b*, a gene previously reported to be downregulated in psoriasis, showed decreased expression, while its mouse ortholog *Defb14* was similarly downregulated in IMQ-induced lesions. Moreover, the UPR-related gene *Hspa5* was consistently induced in lesional skin across both species. Notably, the expression of Gelsolin was significantly reduced in psoriatic lesions of both mice and humans. Pearson correlation analysis showed a negative correlation between *Gelsolin* and *Hspa5* (r = -0.86, *p* = 0.00626) in lesional skin (Figure 8C). Together, these results not only recapitulate known transcriptional features of psoriatic lesions but also highlight a striking and conserved downregulation of Gelsolin, which is closely linked to UPR activation and may play a key role in disease pathogenesis.

**Figure 8.**
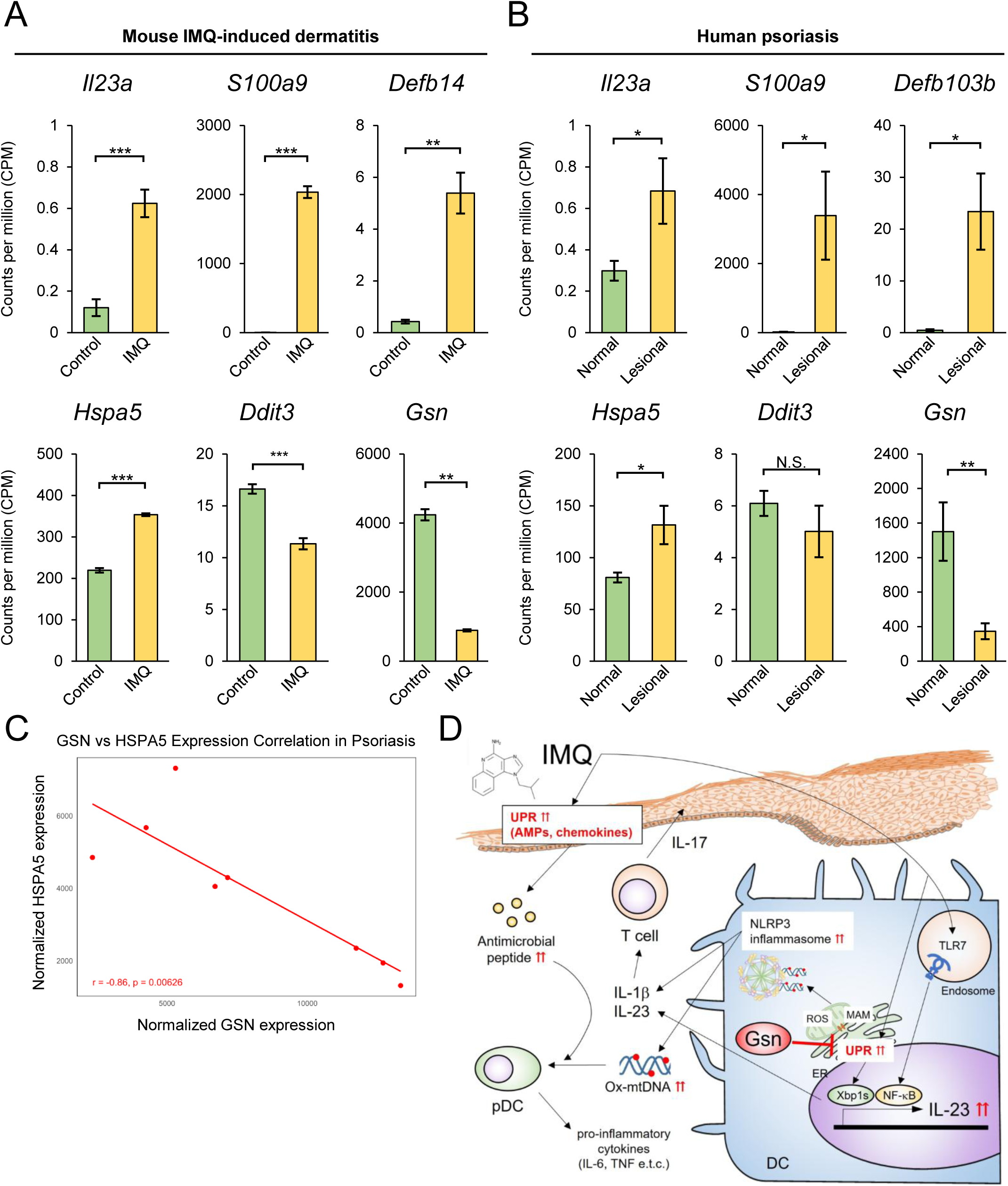
Human and mouse psoriatic lesions exhibit reduced Gelsolin and enhanced ER stress. **a, b,** Datasets (GSE289485 and GSE117405) were re-analyzed, and each gene expression was compared using normalized read counts. (mouse [*n* = 6], human: non-lesionl [*n* = 9], lesional [*n* = 8]) **c,** Pearson correlation analysis between *Gelsolin* and *Hspa5* in (b). d, Schematic representation of this study. IMQ induces ER-mitochondria contact and calcium overload in dendritic cells, triggering UPR activation and mitochondrial dysfunction. Released mtDNA, together with mBD14, activates plasmacytoid DCs, amplifying inflammation. Gelsolin binds IMQ and attenuates these stress responses, with its downregulation in psoriatic lesions correlating with enhanced UPR signaling.

## Discussion

Our study demonstrates that the UPR and the NLRP3 inflammasome exacerbate IMQ-induced psoriasis (Figure 8D). Although the IMQ-induced dermatitis model has provided important insights into IL-23/IL-17-driven skin inflammation, it does not fully recapitulate the immunopathology, chronicity, or therapeutic responsiveness of human plaque psoriasis^44^. Therefore, our findings should be interpreted primarily in the context of IMQ-induced inflammation, and further validation in human psoriasis-relevant systems will be required. Accordingly, phenotypes observed in inflammasome-, TLR7-, or MyD88-dependent settings may reflect mechanisms that are particularly important for IMQ-driven dermatitis rather than universal drivers of human psoriasis. We found that IMQ disrupts mitochondrial homeostasis, leading to excessive ROS production, mitochondrial damage, and the release of mtDNA into the cytoplasm. This is supported by our observation that IMQ-induced IL-1β release from BMDCs was significantly suppressed by the mitochondrial ROS-specific inhibitor mitoquinone (MitoQ), as well as by antioxidants such as GSHEE, NAC, and PDTC. These findings highlight mitochondrial dysfunction as a central driver of inflammasome activation and inflammation in IMQ-induced dermatitis. In addition to its effects on mitochondrial ROS, IMQ increased ER–mitochondria contact sites, a condition commonly associated with facilitated Ca^2+^ dynamics between these organelles (Figure 8D). Although Ca^2+^ transfer can promote mitochondrial ROS production^45,46^, our study does not distinguish which specific Ca^2+^ flux pathways are responsible for the observed effects. Notably, inhibition with 2-APB significantly reduced IMQ-induced ROS generation and IL-1β release; however, because 2-APB modulates multiple Ca^2+^-related processes beyond ER-derived flux, these results should be interpreted as indicating that Ca^2+^ dysregulation in general, rather than a single defined pathway, contributes to IMQ-driven mitochondrial stress and inflammasome activation. While previous studies have shown that IMQ directly targets mitochondrial complex I and NQO2, leading to ROS production and NLRP3 activation^24^, our findings suggest that additional mechanisms, specifically, Ca^2+^ dysregulation and ER-mitochondria interactions, contribute to mitochondrial ROS production.

IMQ also triggered the UPR, characterized by the induction of *Xbp1s* in both DCs and keratinocytes (Figure 8D). Given that ER Ca²⁺ depletion disrupts protein folding and induces ER stress^47^, it is likely that Ca²⁺ release from the ER contributes to UPR activation. Additionally, mitochondrial ROS have been reported to act as upstream signals for ER stress and UPR activation^48,49^, further supporting the link between mitochondrial dysfunction and ER stress responses in IMQ-induced skin inflammation. Our study highlights the critical role of the IRE1α-XBP1s axis in regulating IL-23 expression in DCs. IMQ-induced *Il23a* mRNA expression was partially suppressed by 4μ8c, an IRE1α inhibitor, while overexpression of XBP1s and MyD88 synergistically enhanced the promoter activity of *Il23a* gene, which contains binding sites for both XBP1s and NF-κB. These findings suggest that IL-23 expression in DCs is driven by the convergence of the IRE1α-XBP1s and TLR7-MyD88-NF-κB pathways. Notably, *Il23a* mRNA induction was abolished in MyD88-deficient BMDCs, confirming the indispensable role of TLR7- MyD88 signaling, with XBP1s enhancing its expression. Furthermore, RNA-seq analysis revealed that IMQ failed to induce IL-23 expression in keratinocytes where TLR7 expression is rare but activated the Ca²⁺-IRE1α-XBP1s axis to drive the expression of psoriasis-associated genes. These results suggest that in keratinocytes, IMQ primarily regulates gene expression through the UPR rather than TLR7.

Our data may suggest that IMQ stimulation also promoted the release of oxidized mtDNA into the extracellular environment via NLRP3 inflammasome-dependent pyroptosis (Figure 8D). Previous studies have established that mtDNA acts as a DAMP, activating pathways such as TLR9 and other cytosolic DNA sensors such as cGAS, NLRP3, and AIM2 to drive inflammatory and autoimmune diseases^42,50–53^. Our data further revealed that mtDNA enhanced IL-6 and TNF-α production by pDCs in the presence of mBD14. Additionally, synthetic DNA experiments confirmed that oxidized mtDNA complexed with mBD14 activated pDCs via TLR9, suggesting that mBD14 may protect extracellular mtDNA from DNase-mediated degradation or facilitate its endosomal uptake. Given that *Defb14* expression in keratinocytes is induced via a UPR-dependent mechanism, UPR activation may contribute to the formation of mtDNA-containing immune complexes and associated inflammatory responses under specific inflammatory conditions. However, previous studies have suggested that deficiency of antimicrobial peptides such as CRAMP or β-defensins does not substantially attenuate IMQ-induced dermatitis, indicating that IMQ-driven inflammation can proceed independently of these pathways^12,54,55^.

Beyond immune regulation, metabolic alterations also contribute to psoriasis pathogenesis^40^. High-fat diets have been shown to exacerbate psoriasis via an XBP1- dependent mechanism, promoting mitochondrial ROS production and UPR activation in TLR-stimulated DCs. In our study, IMQ treatment induced UPR activation in psoriatic lesions, and administration of tunicamycin, an ER stress inducer, significantly increased ear thickness and upregulated psoriasis-related gene expression in IMQ-treated mice. Furthermore, carbachol, an IP_3_R activator known to trigger UPR and NLRP3 inflammasome activation, exacerbated IMQ-induced dermatitis and enhanced the expression of psoriasis-related genes. These findings reinforce the role of the Ca²⁺-UPR axis in psoriasis pathology. Importantly, 2-APB significantly suppressed caspase-1 activation in BMDCs and reduced IL-17 expression in skin lesions, ultimately ameliorating psoriasis symptoms in vivo. Collectively, these results suggest that Ca²⁺ transfer from the ER to mitochondria, followed by ER stress and NLRP3 inflammasome activation, represents an important pathogenic mechanism in IMQ-induced dermatitis. Whether this pathway plays an equivalent role in human psoriasis remains to be established. Although IL-1 family cytokines and inflammasome activation contribute to inflammation in several experimental models of psoriasis-like dermatitis, clinical studies targeting IL-1 signaling have not demonstrated therapeutic efficacy comparable to IL-23- or IL-17-directed therapies in plaque psoriasis^40,44^. Therefore, the contribution of NLRP3 inflammasome activation identified in this study should be interpreted primarily within the context of IMQ-induced inflammation. In addition, the vehicle formulation of Aldara cream was not commercially available to us, precluding inclusion of an Aldara vehicle control in the present study. Although this represents a limitation of the in vivo experiments, our in vitro findings obtained using purified IMQ support the conclusion that the observed inflammatory responses can be induced by IMQ itself. Nevertheless, further studies using appropriate vehicle controls will be valuable to more precisely define the contribution of the cream formulation.

We identified Gelsolin as a critical regulator that suppresses IMQ-induced ER stress responses (Figure 8D). Importantly, the rationale for focusing on Gelsolin is supported not only by its interaction with IMQ but also by independent observations demonstrating altered Gelsolin expression in human psoriasis^37,56^. Gelsolin-deficient cells exhibited enhanced activation of UPR upon IMQ stimulation. These findings are consistent with a recent study in macrophage-specific Gelsolin-deficient mice, which demonstrated increased mitochondria damage and NLRP3 inflammasome-mediated IL- 1β secretion and aggravated arthritis, indicating a broader role for Gelsolin in modulating inflammation^36^. Given its established function in actin remodeling^33,34^, Gelsolin may influence the structural integrity of ER-mitochondria contact sites, which are essential for Ca^2+^ exchange between the two organelles. The loss of Gelsolin could increase these contact sites, resulting in excessive Ca^2+^ influx into mitochondria and subsequent mitochondrial dysfunction, including ROS generation. Additionally, Gelsolin’s ability to bind cytosolic Ca^2+^ may contribute to its regulatory function. By buffering excess Ca^2+^,

Gelsolin may prevent aberrant Ca^2+^ signaling that would otherwise promote ER stress and inflammasome activation. These findings highlight Gelsolin as a key modulator of intracellular Ca^2+^ dynamics, ER stress responses, and inflammatory signaling pathways, offering new insights into its potential as a therapeutic target in inflammatory skin diseases such as psoriasis.

Importantly, our analysis of human psoriatic lesions revealed significant Gelsolin downregulation alongside upregulation of ER stress genes such as *Hspa5* (Figure 9B). Furthermore, Pearson correlation analysis showed a negative correlation between *Gelsolin* and *Hspa5*, suggesting that reduced *Gelsolin* expression exacerbates ER stress and inflammation in human psoriasis. These findings in human psoriasis patients support a protective role for Gelsolin and suggest that its downregulation in psoriatic lesions may contribute to disease-associated inflammatory responses. Gelsolin supplementation or strategies to enhance its expression could thus represent novel therapeutic approaches.

Our findings reveal a previously unrecognized interplay between the NLRP3 inflammasome, ER stress, and Ca^2+^ homeostasis in IMQ-induced psoriasis (Figure 8D). Furthermore, IMQ-induced mtDNA release amplifies inflammation through pDC activation in the presence of antimicrobial peptides, such as mBD14, while Gelsolin serves as a protective factor by regulating ER stress and mitochondrial function. These findings suggest that targeting ER-mitochondria interactions, and UPR-associated molecules could offer novel therapeutic strategies for psoriasis. Accordingly, the mechanistic pathways identified in this study should be interpreted primarily as regulators of IMQ-induced inflammation, and their contribution to human psoriasis will require validation in disease-relevant experimental systems and clinical samples.

## Material and Methods

### Mice

C57BL/6J mice were purchased from CLEA Japan (Tokyo, Japan) and maintained in accordance with the guidelines of the Committee on Animal Research at the Nara Institute of Science and Technology. MyD88-deficient mice were purchased from Oriental Bio Service.

### Cells

HeLa cells, HEK293T cells, and MEFs were cultured in Dulbecco’s modified Eagle’s medium (DMEM) supplemented with 10% fetal bovine serum (FBS) at 37°C in a humidified 5% CO_2_ incubator. Mouse BMDCs and pDCs were obtained by culturing mouse bone marrow cells in RPMI 1640 medium (Nacalai Tesque) supplemented with 10% FBS, 1% antibiotic-antimycotic mixed stock solution (Nacalai Tesque), 0.1 mM 2- mercaptoethanol, and 10 ng/mL recombinant mouse GM-CSF (PeproTech) or 100 ng/mL mouse Flt-3 ligand (BioLegend) for 6–8 days.

### Isolation and culture of primary mouse keratinocytes

Primary mouse keratinocytes were isolated from neonatal mouse skin. Briefly, peeled mouse skin was washed with PBS and then incubated with CnT-Prime medium (CELLnTEC) containing 2.5 mg/mL dispase II and 1% penicillin-streptomycin mixed solution (Nacalai Tesque) at 4°C overnight. The next day, the skin was washed with PBS and the epidermis was isolated from the dermis. The epidermis was incubated with accutase (Nacalai Tesque) for 20 minutes at room temperature to isolate keratinocytes. Keratinocytes were passed through a 100-μm filter and pelleted. Keratinocytes were resuspended in CnT-Prime medium containing 1% penicillin-streptomycin mixed solution and seeded onto culture dishes precoated with Cellmatrix Type I-C (Nitta Gelatin).

### Reagents and plasmids

All ligands for pattern recognition receptors were purchased from Invivogen unless otherwise stated. LPS and ATP were purchased from Sigma-Aldrich. Working concentrations of each innate immune ligand were as follows: LPS (20–1000 ng/mL), imiquimod (5–50 μg/mL), resiquimod (50 μg/mL), ATP (5 mM), nigericin (5 μM), and D19 (100 nM). Recombinant mouse beta-defensin 14 was purchased from Prospec and used at 5 μg/mL. Each inhibitor was used at the following concentrations: PDTC (10 μM, Sigma-Aldrich), GSHEE (10 mM, Sigma-Aldrich), NAC (10 or 20 mM, Thermo Fisher Scientific), 2-APB (50 μM, Wako), 4μ8c (25 or 32 μM, Cayman), mitoquinone (10 μM, Wako). Inhibitors were added directly to the cell culture medium 0.5–1 h before stimulation. Tunicamycin (LKT-Labs) and thapsigargin (Wako) were used to induce ER stress.

To overexpress Xbp1s, the mouse Xbp1s coding sequence (CDS) was inserted into a pFLAG-CMV2 expression vector (Sigma-Aldrich). For the reporter assay, the mouse *Il23a* genomic fragment (−2,030 to +20) was amplified by PCR and inserted into the pGL3-promoter (Promega).

### Synthesis of oxidized DNA fragment

Oxidized DNAs were generated by PCR in the presence of 200 μM 8-Oxo-2′-dGTP (TriLink) and then purified using a QIAquick Nucleotide Removal Kit (Qiagen). Synthesized oxidized DNA fragments and mBD14 were mixed in FBS-free RPMI 1640 and incubated for 15 min at room temperature prior to stimulation. The primers used for oxidized DNA synthesis are described in Supplementary Table 10.

### Flow cytometry-based cell death assay

LPS primed or unprimed BMDCs were seeded onto uncoated 96-well plates at a density of 1.0 × 10^5^ cells per well. After 3 or 18 h of stimulation, cells were harvested and stained with 200 ng/mL propidium iodide (PI, Nacalai Tesque) and Annexin V-APC (BD Bioscience) in Annexin V binding buffer (10 mM Hepes-NaOH [pH 7.2], 140 mM NaCl, 2.5 mM CaCl_2_). Data were collected by FACS Accuri (BD Bioscience).

### Measurement of intracellular ROS

BMDCs were seeded onto untreated 96-well plates at a density of 1.0 × 10^5^ cells per well. After 30 min of stimulation with 50 μg/ml IMQ, CellROX™ Deep Red (Thermo Fisher Scientific) was added directly to the culture medium at a final concentration of 1 μM and incubated at 37°C for 15 min. The cells were then washed with PBS and the fluorescence intensity was analyzed by FACS Accuri. To measure mitochondrial ROS, 5 × 10^5^ BMDCs were seeded onto untreated 24-well plates and incubated with 2.5 μM MitoSOX™ Red Mitochondrial Superoxide Indicator (Thermo Fisher Scientific) in RPMI 1640 medium for 30 min. The cells were then washed with RPMI 1640 medium and seeded onto untreated 24-well plates followed by stimulation with 50 μg/ml IMQ. After 30 min of IMQ stimulation, the cells were washed with PBS and analyzed by flow cytometry.

### PLA

BMDCs were seeded at a density of 1.5 × 10^5^ onto 24-well plates with poly-L-lysin- coated coverslips and stimulated with LPS for 3 h followed by stimulation with 50 μg/mL IMQ. The cells were then washed with PBS, fixed in 10% formaldehyde, and incubated at room temperature. After 10 min, fixation was stopped using 1 M glycine and the cells were washed with PBS. The coverslips containing the cells were stored in 100 mM glycine. The cells were permeabilized in PBS containing 0.1% Triton-X100 for 20 min at room temperature and then washed twice with PBS. Blocking solution was then added and the cells were incubated at 37°C for 30 min. The cells were then treated with PBS containing anti-VDAC1/3 antibody (ab14734, Abcam) and anti-IP_3_R1 antibody (an264281, Abcam) at 4°C overnight and washed three times with TBS-T. Subsequent experiments were performed using kits (Duolink In Situ PLA Probe Anti-Mouse MINUS [DUO92004, Sigma-Aldrich] and Duolink In Situ PLA Probe Anti-Rabbit PLUS [DUO92002, Sigma-Aldrich]) according to the manufacturer’s instructions. Nuclei were stained with Hoechst33342. PLA signals were visualized by fluorescence microscopy and the number of PLA dots per cell was counted.

### Mitochondrial Ca^2+^ measurement

First, 5 × 10^5^ BMDCs were seeded onto untreated 24-well plates and incubated with 5 μM Rhod-2-AM (Dojindo) in RPMI 1640 for 30 min. The cells were then washed three times with HBSS and seeded onto untreated 24-well plates, followed by stimulation with 50 μg/ml IMQ. After 30 min of stimulation, the cells were washed with HBSS and analyzed by FACS Accuri.

### Mitochondrial membrane potential analysis

BMDCs were seeded onto untreated 96-well plates at a density of 1.0 × 10^5^ cells per well. After 1 h of stimulation with 50 μg/ml IMQ, the cells were stained with 2 μM JC-1 (Dojindo) for 30 min at 37°C in RPMI 1640 medium. The cells were washed with PBS and resuspended with imaging buffer (Dojindo) for analysis by FACS Accuri.

### Cellular fractionation and measurement of mtDNA

Stimulated cells were washed with PBS and lysed with digitonin lysis buffer (50 mM HEPES-NaOH [pH 7.4], 150 mM NaCl, 20 μg/ml digitonin). The cell lysate was centrifuged at 1000 ×*g* for 3 min at 4°C, and the supernatant was collected into a new tube. This step was repeated three times to remove intact cells and cell debris. The supernatant was then centrifuged at 17,000 ×*g* for 10 min at 4°C. The supernatant was collected as the cytoplasmic fraction and the pellet was collected as the mitochondrial fraction. DNA from the cytosolic fraction and cell culture supernatant were purified using the QIAquick Nucleotide Removal Kit (Qiagen). Pellets containing mitochondria were lysed with cell lysis buffer (100 mM Tris-HCl [pH 8.0], 5 mM EDTA [pH 8.0], 0.2% SDS, 200 mM NaCl) containing 1/100 volume of protease K at 56°C for 2 h. DNA from mitochondrial lysates was precipitated with isopropanol and washed with 70% ethanol. The amount of mitochondrial DNA in each fraction was analyzed by qPCR using PowerUP SYBR Green PCR Master Mix (Applied Biosystems) with a Step One Real-Time PCR System (Applied Biosystems).

### ELISA

BMDCs were seeded onto 96-well plates at a density of 1.0 × 10^5^ cells per well. For IL- 1β measurements, cells were preincubated with 20 ng/ml of LPS for 3 h and stimulated with 50 μg/ml IMQ, 50 μg/ml RSQ, 5 mM ATP, or 5 μM nigericin for 30 min or 3 h. In the case of pDCs, the cells were seeded onto 96-well plates at a density of 2.0 × 10^5^ cells per well and stimulated with ligands for 24 h. The cytokine levels of IL-6, TNF-α, and IL-1β in the culture supernatant were measured using a mouse IL-6 DuoSet ELISA (R&D Systems), mouse TNF-α ELISA kit (Invitrogen), and mouse IL-1β ELISA kit (Invitrogen), respectively, according to the manufacturer’s instructions.

### Western blot analysis

BMDCs were seeded onto six-well plates at a density of 1.0 × 10^6^ cells per well. For caspase-1, the cells were pretreated with 20 ng/ml LPS for 3 h and stimulated with 50 μg/ml IMQ for 30 min. The cells were washed with PBS and lysed with 1× SDS sample buffer (25 mM Tris-HCl [pH 6.8], 5% glycerol, 1% SDS, 50 mM DTT, and BPB). For PERK and eIF2α, cells were stimulated with IMQ, RSQ, or 1 mM DTT for 0.5–6 h and lysed with lysis buffer (150 mM NaCl, 1% Nonidet P-40, 1 mM EDTA, 20 mM Tris-HCl [pH 7.5]) containing cOmplete™ Protease Inhibitor Cocktail (Roche). The cell lysates were subjected to SDS-PAGE and transferred to an Immun-Blot PVDF membrane (Bio-Rad). The membrane was then immunoblotted with anti-caspase-1 (p20) (AG-20B-0042- C100, AdipoGen), anti-phospho-PERK (Thr980) (#3179, CST), anti-PERK (#3192, CST), anti-phospho-eIF2α (Ser51) (#3398, CST), or anti-eIF2α (#9722, CST) antibodies. Horseradish-peroxidase-conjugated anti-mouse, rabbit, or goat IgG (Sigma-Aldrich) were used as secondary antibodies. Protein bands were visualized using Western Lightning Plus-ECL (PerkinElmer) and analyzed on a LAS-4000 (Fujitsu Life Sciences). Image files were processed using ImageJ software 1.8.0_172 (NIH).

### Ca^2+^ flux analysis

BMDCs or keratinocytes were incubated with Quest Fluo-8 (ABD Bioquest) for 30 min in the dark at room temperature. The cells were washed three times with HBSS containing 1 mM MgCl_2_, 2 mM CaCl_2_, and 1% FBS, and the Ca^2+^ flux was measured after stimulation with 50 μg/ml IMQ or 50 μg/ml RSQ. Fluo-8 intensity was monitored every 5 seconds on an LSM700 or LSM710 (Zeiss). Keratinocytes were washed with CnT-PR and the Ca^2+^ flux was measured. Ionomycin was used as a positive control. Ca^2+^- insensitive fluorescence was subtracted from each wavelength before calculations to normalize fluorescence values. The values were then plotted against time and shown as F/F0.

### RNA isolation, reverse transcription PCR, and quantitative real-time PCR

Total RNA was isolated using Trizol reagent (Invitrogen) and cDNA was synthesized using ReverTra Ace (Toyobo) according to the manufacturer’s instructions. To amplify spliced and unspliced Xbp1 for RT-PCR, Go Taq Master Mix (Promega) was used with the primers described in Supplementary Table 11. PCR conditions were 2 min at 95°C, 35 cycles of denaturation at 95°C for 30 sec, annealing at 60°C for 30 sec, and extension at 72°C for 2 min. A final extension was performed at 72°C for 7 min. PCR products were electrophoresed on 2.5% agarose, and spliced and unspliced Xbp1 bands were visualized using Midori Green Advance DNA Stain (Nippon Genetics). Real-time quantitative PCR (RT-qPCR) was performed using PowerUP SYBR Green PCR Master Mix with Step One. The primers used for RT-qPCR are described in Supplementary Table 12. Relative gene expression was calculated using the comparative Ct (2^−ΔΔCt^) method. Expression levels were normalized to 18S rRNA, Actb, or Gapdh, depending on the experiment, and are presented relative to the corresponding untreated control.

### Luciferase reporter assay

To measure *Il23a* promoter activity, 7.5 × 10^5^ HEK293T cells were seeded onto 24-well plates and transfected with 100 ng of pGL3-Il-23p19 and 500 ng of expression plasmid. As an internal control, 10 ng of pRL-TK (Promega) was transfected simultaneously. The medium was changed 6 h after transfection. After 24 h of transfection, luciferase activity was measured using a TriStar2 LB 942 Multidetection Microplate Reader (Berthold) with the Dual-Glo Luciferase System (Promega) according to the manufacturer’s instructions. To measure UPR activity, transfected HeLa cells were stimulated with 50 μg/ml IMQ or 50 μg/ml RSQ for 9 h and the luciferase activity was analyzed.

### Induction of IMQ-induced psoriasis-like skin inflammation

Mice aged 8–12 weeks were treated with 10 μg tunicamycin or 20 μg 2-APB applied to the ears or injected intraperitoneally with 25 μg carbachol, followed by the application of 12.6 mg IMQ cream (5%) (Mochida Pharmaceutical) to the ears, for 6–7 consecutive days. As a control, DMSO or EtOH was applied to the opposite ear. For histological analysis, approximately the distal 2 mm of each ear (outer tip region) was excised using the same anatomical landmarks for all mice to ensure standardized tissue sampling. Ear samples were fixed in 4% paraformaldehyde phosphate buffer overnight at 4°C and then embedded in paraffin. Sections were cut at 5 μm and stained with hematoxylin and eosin.

### Identification of IMQ-binding proteins

#### [Immobilization of IMQ onto magnetic beads]

Carboxyl-functionalized magnetic beads (2.5 mg; Tamagawa Seiki Co., Ltd.) were washed several times with N,N-dimethylformamide (DMF) and activated with N-hydroxysuccinimide (NHS, 1 M in DMF) and 1-ethyl-3-(3-dimethylaminopropyl) carbodiimide hydrochloride (EDC·HCl). After incubation at room temperature for 2 h, the beads were washed with DMF and resuspended to obtain NHS-activated beads. For ligand coupling, NHS-activated beads (1.25 mg per reaction) were incubated with a ligand solution containing 2 mM IMQ and 4 mM triethylamine in DMF for 70 min at room temperature with gentle mixing. Control beads were treated in parallel with DMF alone. Following the coupling reaction, residual NHS groups were quenched with 1 M ethanolamine for 2 h. Finally, the beads were washed several times with 50% methanol and resuspended in 50% methanol for storage until use.

#### [Preparation of BMDC-derived protein lysates]

BMDCs (2.0 × 10⁶ cells) were collected, washed once with PBS, and pelleted again. The cell pellets were resuspended in KCl buffer (150 mM KCl, 0.5% Nonidet P-40, 20 mM HEPES [pH 7.9], 1 mM MgCl₂, 0.2 mM CaCl₂, 0.2 mM EDTA, 1 mM DTT, supplemented with protease inhibitor cocktail). After incubation on ice for 5 min, lysates were clarified by centrifugation at 20,000 ×*g* for 5 min at 4°C, and the supernatants were collected as protein extracts.

#### [Purification of IMQ-binding proteins]

IMQ-conjugated beads were washed with KCl buffer and separated using a magnetic rack. Protein lysates prepared as described above were added to the beads, and the mixture was incubated at 4°C for 4 h with gentle rotation. After incubation, the beads were washed five times with KCl buffer without DTT. Bound proteins were eluted by adding 3× SDS sample buffer containing 15% 2-mercaptoethanol and protease inhibitors, followed by heating at 95°C for 5 min. The supernatants containing eluted proteins were collected after cooling on ice.

#### [SDS-PAGE and silver staining]

Proteins eluted from the beads were separated on a 12% polyacrylamide gel prepared using the TGX FastCast Acrylamide Kit (Bio-Rad) according to the manufacturer’s instructions. Samples were loaded onto the gel and electrophoresed in running buffer (0.25 M Tris, 192 mM glycine, 1% SDS) at 20 mA for 60 min. Separated proteins were visualized by silver staining using the Silver Stain MS Kit (Wako) and subsequently subjected to mass spectrometry analysis.

### Pulldown assay

HEK293T cells were seeded onto 6-well plates (Falcon) coated with Cellmatrix Type I-C (Nitta Gelatin) at 8.0 × 10⁵ cells per well. Cells were transfected with FLAG-Gelsolin expression plasmid using Lipofectamine 2000 (Invitrogen) according to the manufacturer’s protocol. Two days post-transfection, cells were washed with PBS, and lysed in 150 μL of lysis buffer (20 mM HEPES-HCl [pH 7.5], 150 mM NaCl, 1% NP-40, 1 mM EDTA) supplemented with protease inhibitors. Lysates were incubated on ice for 5 min and clarified by centrifugation at 20,000 ×*g* for 5 min at 4°C. The supernatants were collected as cell lysates. Cell lysates were incubated with IMQ-conjugated beads (described before) overnight at 4°C with gentle rotation. After incubation, the bead-bound proteins were eluted, separated by SDS-PAGE, subjected western blot analysis for FLAG- Gelsolin and β-Actin.

### Immunofluorescence

WT and Gsn-deficient MEFs were seeded and transfected with pLZR-c-FLAG-empty or pLZR-c-FLAG-mGelsolin plasmid, and cultured for 24 h. For immunofluorescence, cells were then replated onto poly-L-lysine–coated coverslips (Matsunami) in 24-well plates and cultured overnight. Cells were stimulated with IMQ (50 μg/mL) for 0-15 min, washed with PBS, and fixed with 4% paraformaldehyde for 20 min. After permeabilization and blocking, cells were incubated with primary antibodies overnight at 4°C: anti-FLAG Ab (SigmaAldrich), anti-Calnexin Ab (Abcam), and anti-TOM20 Ab (CST). After washing, cells were incubated with Alexa Fluor–conjugated secondary antibodies (Invitrogen) for 1 h and counterstained with Hoechst 33342 (Dojindo). Images were acquired on an LSM900 confocal microscope (Zeiss) using 405, 488, and 561 nm lasers, and processed with ZEN software.

### Generation of Gelsolin-deficient MEFs

MEFs were electroporated with a px330 plasmid expressing Cas9 and a single-guide RNA targeting the Gelsolin coding sequence (Table S13). Electroporation was performed using Neon Transfection System (Thermo Fisher) according to the manufacturer’s instructions. After recovery, cells were cultured under standard conditions, and single cells were sorted by flow cytometry into 96-well plates to establish clonal populations. Genomic DNA was extracted from individual clones, and the target locus was amplified by PCR and analyzed by Sanger sequencing to confirm the presence of indels and determine the genotype (Figure S11).

### RNA sequencing

Keratinocytes were stimulated with 50 μg/ml IMQ in the presence or absence of 25 μM 4μ8c. After 3 h of stimulation, total RNA was isolated from keratinocytes using the FastGene™ RNA Premium Kit (Nippon Genetics) and RNA quality was analyzed using the Agilent 2100 Bioanalyzer. Illumina next-generation sequencing analysis was performed by Genewiz (Tokyo, Japan). Raw sequencing reads from both in-house and public RNA-seq datasets were quality-trimmed using Trim Galore and aligned to the appropriate reference genome using HISAT2. Alignment files were sorted using SAMtools, and reads mapping to rRNA regions were removed using BEDTools. Gene-level read counts were generated using featureCounts (Rsubread package) with GENCODE gene annotations. Gene annotations, including gene symbols and gene biotypes, were assigned using GENCODE annotation files and the rtracklayer package. Pseudogenes were excluded from downstream analyses.

For in-house RNA-seq data, lowly expressed genes were filtered using filterByExpr, and library sizes were normalized using the trimmed mean of M-values (TMM) method implemented in edgeR. Differential gene expression analysis was performed using the exactTest function in edgeR.

Public RNA-seq datasets were obtained from the Gene Expression Omnibus (GEO) database. GSE289485 was selected because it contains transcriptomic profiles from IMQ-induced psoriasis-like dermatitis in mice, allowing comparison with IMQ-responsive transcriptional changes observed in our keratinocyte model. GSE117405 was selected because it contains transcriptomic profiles from lesional skin of patients with psoriasis, enabling assessment of the overlap with human psoriasis-associated inflammatory signatures. Detailed information on the datasets and analyzed samples is provided in Supplementary Table 14. Public RNA-seq data were analyzed using DESeq2. Genes with very low expression levels (total read count < 6 across all samples) were excluded, and differential expression analysis was performed using the Wald test with multiple-testing correction by the Benjamini–Hochberg method.

### Identification of overlapping inflammatory transcriptional signatures associated with mouse IMQ-induced dermatitis and human psoriasis

To identify overlapping inflammatory transcriptional signatures associated with mouse IMQ-induced dermatitis and human psoriasis, genes responsive to IMQ stimulation and upregulated in mouse IMQ-treated skin were extracted using the following criteria: log2 fold change (FC) ≥ 1 and *p* value < 0.05. After removal of duplicate gene entries, 286 mouse genes were identified. Mouse genes were converted to their corresponding human orthologs using BioMart. Genes without human orthologs were excluded, and only genes with a human orthology confidence score of 1 were retained. After removal of duplicate entries based on human gene symbols, a total of 238 human orthologous genes were obtained.

Separately, genes upregulated in human psoriasis skin samples were identified from the public RNA-seq dataset (GSE117405) using the criteria of log2FC ≥ 1 and adjusted *p* value (*padj*) < 0.05. After removal of duplicate gene entries, 2,192 genes upregulated in psoriasis skin samples were identified. Overlapping genes between the mouse-derived human orthologous gene set and the human psoriasis upregulated gene set were identified, resulting in 38 overlapping upregulated genes. The complete list of overlapping genes is provided in Supplementary Table 4.

### Statistics

Statistical significance was determined using the Student’s *t-*test and ANOVA with Tukey’s test. *P* < 0.05 was considered statistically significant.

### Study approval

All murine experimental protocols were approved by the Animal experimental Committee at the Nara Institute of Science and Technology (the approval no.2310).

## Data availability

All data supporting the findings of this study and its supplementary information files are available within the article.

## Acknowledgments

We thank C. Suzuki for secretarial assistance, and K. Takahashi and A. Takara for technical support. This work was supported by the Japan Society for the Promotion of Science KAKENHI (grant numbers 17H04066, 19K22533, 20H03468, 23K27392, and 24K22057 to T. Kawai; 17K15726, and 21K14817 to D. Ori; 19K07608 to T. Kawasaki; 23K14546 to N. Kano). This work was also supported by ONO PHARMACEUTICAL CO., LTD., the Takeda Scientific Foundation to T. Kawai and K. Kohno, and Japan Agency for Medical Research and Development (grant number 223fa727001h0001).

## Author contributions

DO, HO, TKawasaki, KJI, KN, MY, KKohno, and TKawai designed the experiments. DO, HO, RK, MM, SH, ST, TT, RT, and TKawasaki performed the experiments. KKobiyama, HN, MS, YK, and AT prepared the materials. DO, HO, and TKawai wrote the manuscript. All authors edited and approved the final draft. DO was listed first for having a larger role in performing experiments and analyzing data.

## Disclosure and competing interests statement

The authors declare that they have no conflict of interest.

---

**Supplemental Figure 1.**
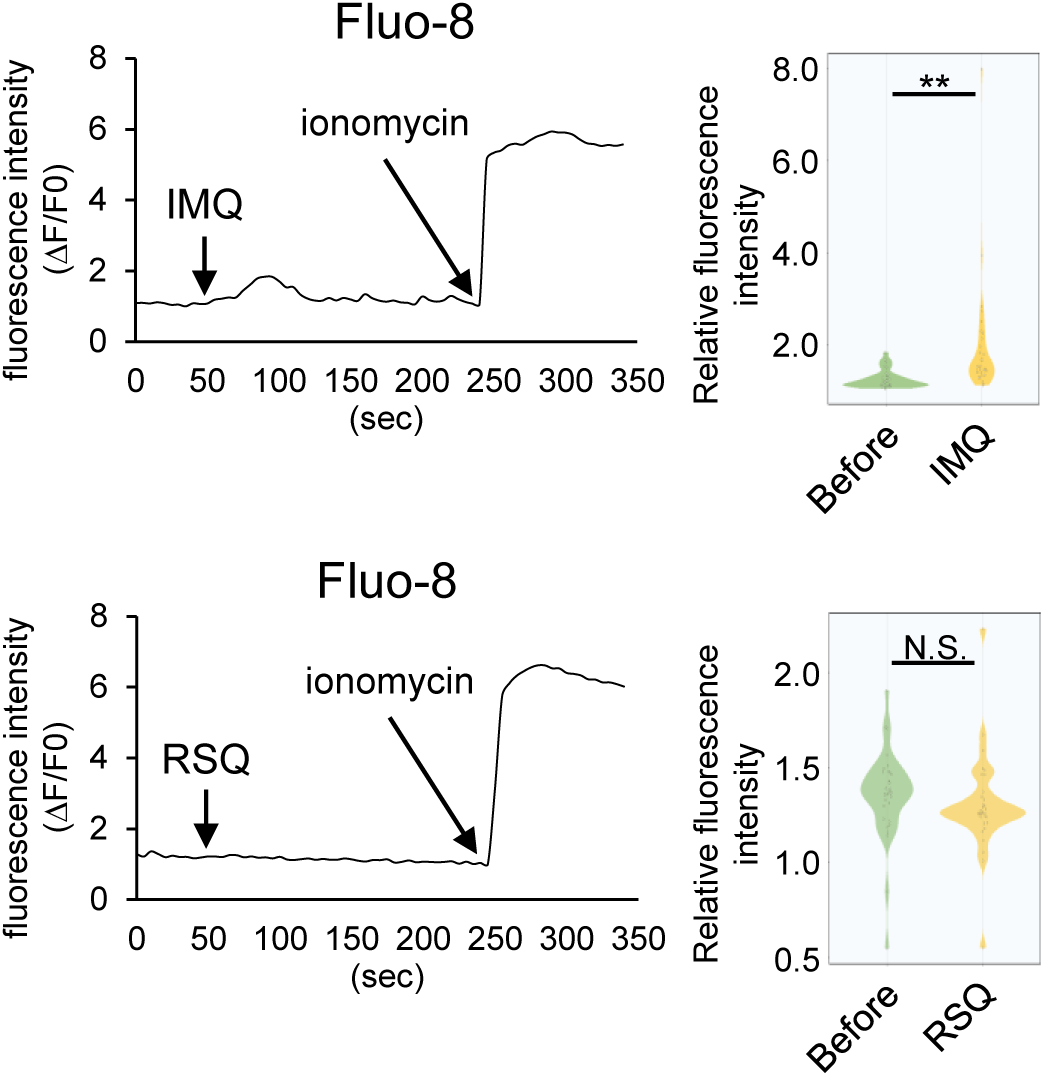
IMQ induces the formation of ER-mitochondria contact sites to transfer Ca^2+^ from the ER to mitochondria for NLRP3 inflammasome activation. BMDCs were pre-stained with Fluo-8 followed by IMQ or RSQ stimulation. Ca^2+^ influx was monitored by fluorescent microscopy. For analysis, fluorescence was normalized (ΔF/F0). Left panel shows representative Fluo-8 fluorescent intensity images. Right panel shows violin plots (IMQ: *n* = 20). Data are presented as the mean ± s.e.m. Each circle indicates an independent biological sample. *P*-values were calculated using Student’s *t*-test. (N.S.: not significant, **: *p* < 0.01).

**Supplemental Figure 2.**
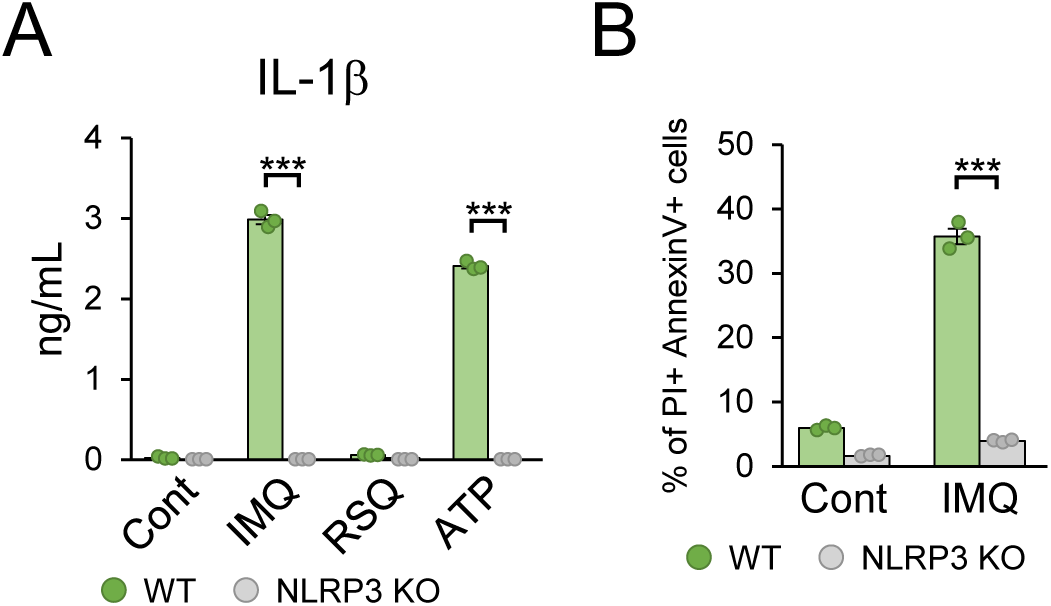
IMQ activates NLRP3 inflammasome. WT and NLRP3-deficient BMDCs were pretreated with LPS, followed by stimulation with the indicated activators and IL-1β concentrations in cell culture supernatants were measured by ELISA (n = 3) (a). Cells were stained with PI and Annexin V, and cell death was analyzed by flow cytometry (n = 3) (b). Data are presented as the mean ±s.e.m. Each circle indicates an independent biological sample. *P*-values were calculated by one-way ANOVA with Tukey’s test. (***: *p* < 0.001).

**Supplemental Figure 3.**
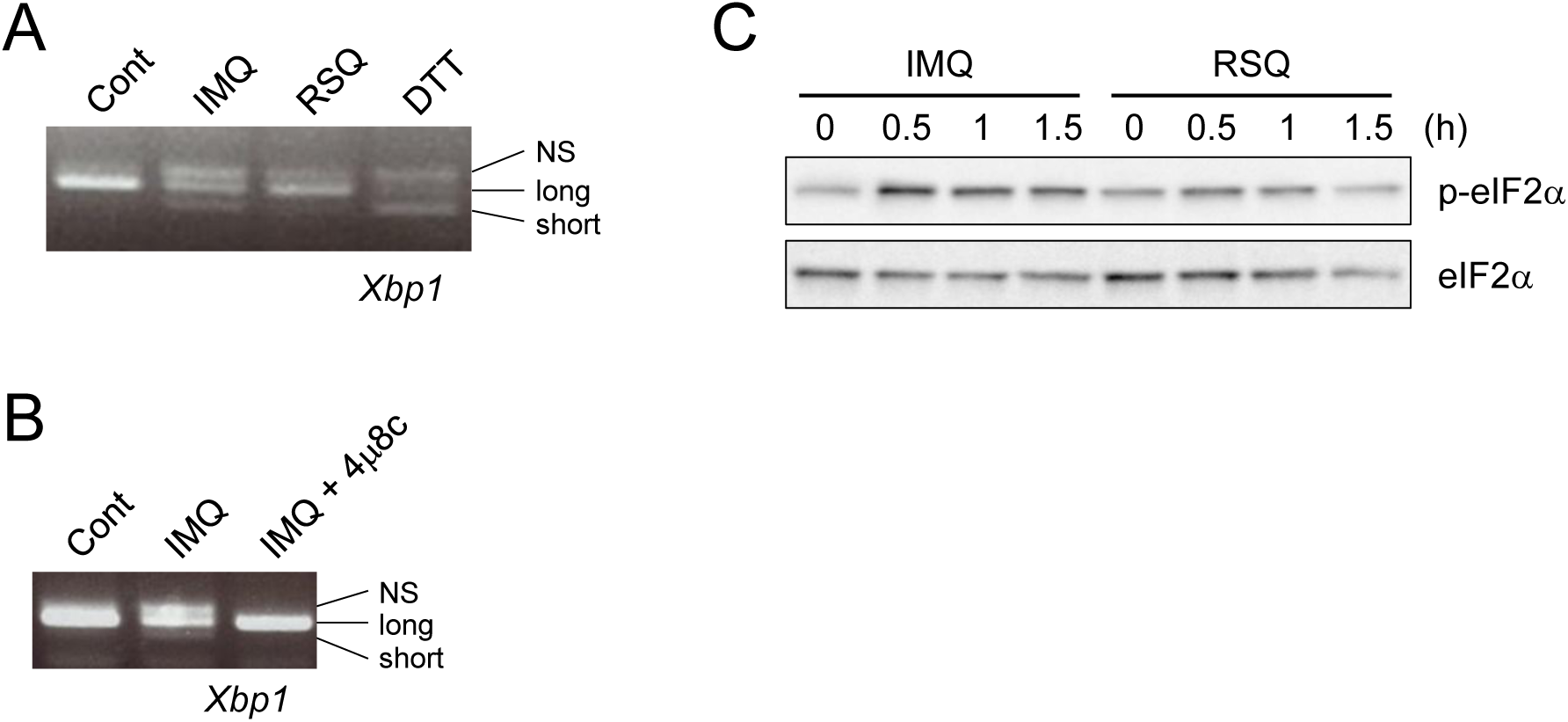
IMQ promotes UPRs in DCs. **(A)** BMDCs were stimulated with IMQ, RSQ, or DTT. Total RNA was harvested and *Xbp1s* mRNA was amplified by PCR. Data are shown as a representative gel electrophoresis image. **(B)** BMDCs pretreated with 4μ8c followed by IMQ stimulation. Total RNA was harvested and *Xbp1s* mRNA was amplified by PCR. Data are shown as a representative gel electrophoresis image. **(C)** BMDCs were stimulated with IMQ or RSQ. Cell lysates were analyzed by immunoblotting for the phosphorylated form of eIF2α. Data are shown as a representative immunoblot analysis images.

**Supplemental Figure 4.**
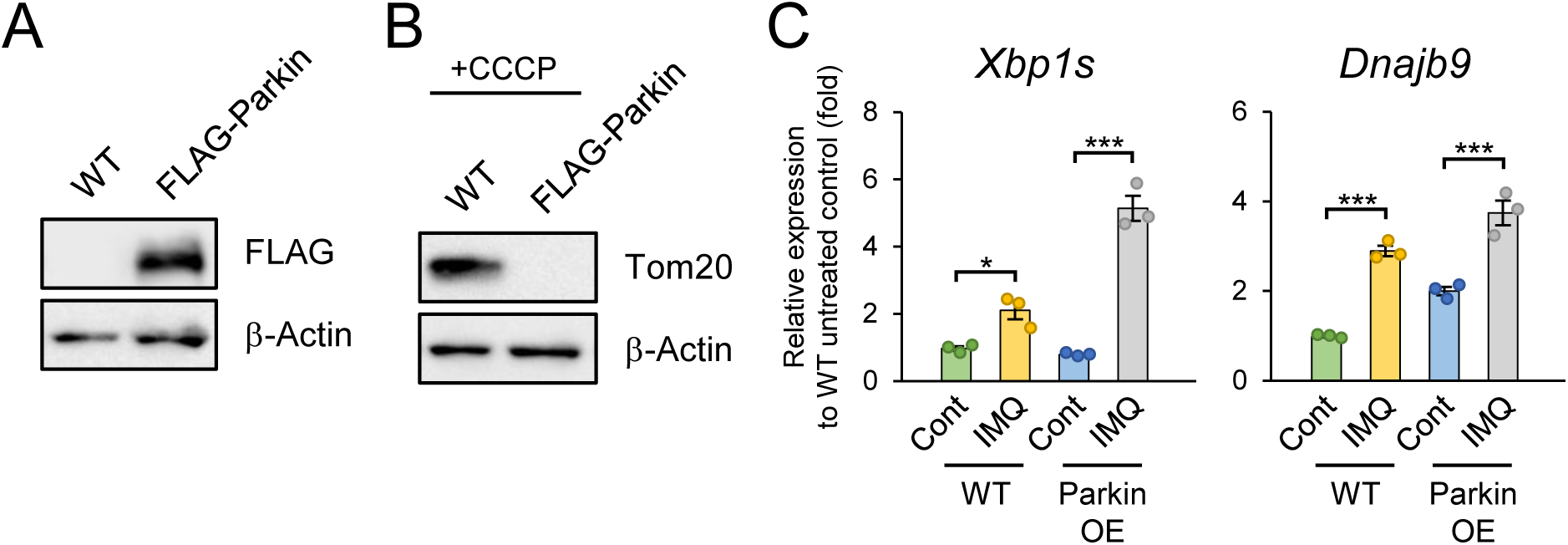
Mitochondria is dispensable to IMQ-induced UPRs in DCs. (A) Cell lysates from WT and Perkin-expressing RAW264.7 cells were analyzed by immunoblotting for Parkin expression. Data are shown as a representative immunoblot analysis images. (B) WT and Perkin-expressing RAW264.7 cells were treated with CCCP. Cell lysates were analyzed by immunoblotting for Tom20 expression. Data are shown as a representative immunoblot analysis images. (C) WT and Perkin-expressing RAW264.7 cells were treated with CCCP, followed by IMQ stimulation, and the expression of short form Xbp1 (Xbp1s) and Dnajb9 (Erdj4) mRNA was measured by qPCR (n = 3). Data are presented as the mean ± s.e.m. Each circle indicates an independent biological sample. *P*-values were calculated by one-way ANOVA with Tukey’s test (C). (*: *p* < 0.05, ***: *p* < 0.001).

**Supplemental Figure 5.**
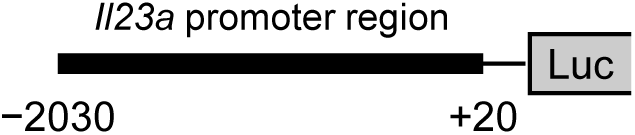
Schematic diagram of the plasmid used in the luciferase assay.

**Supplemental Figure 6.**
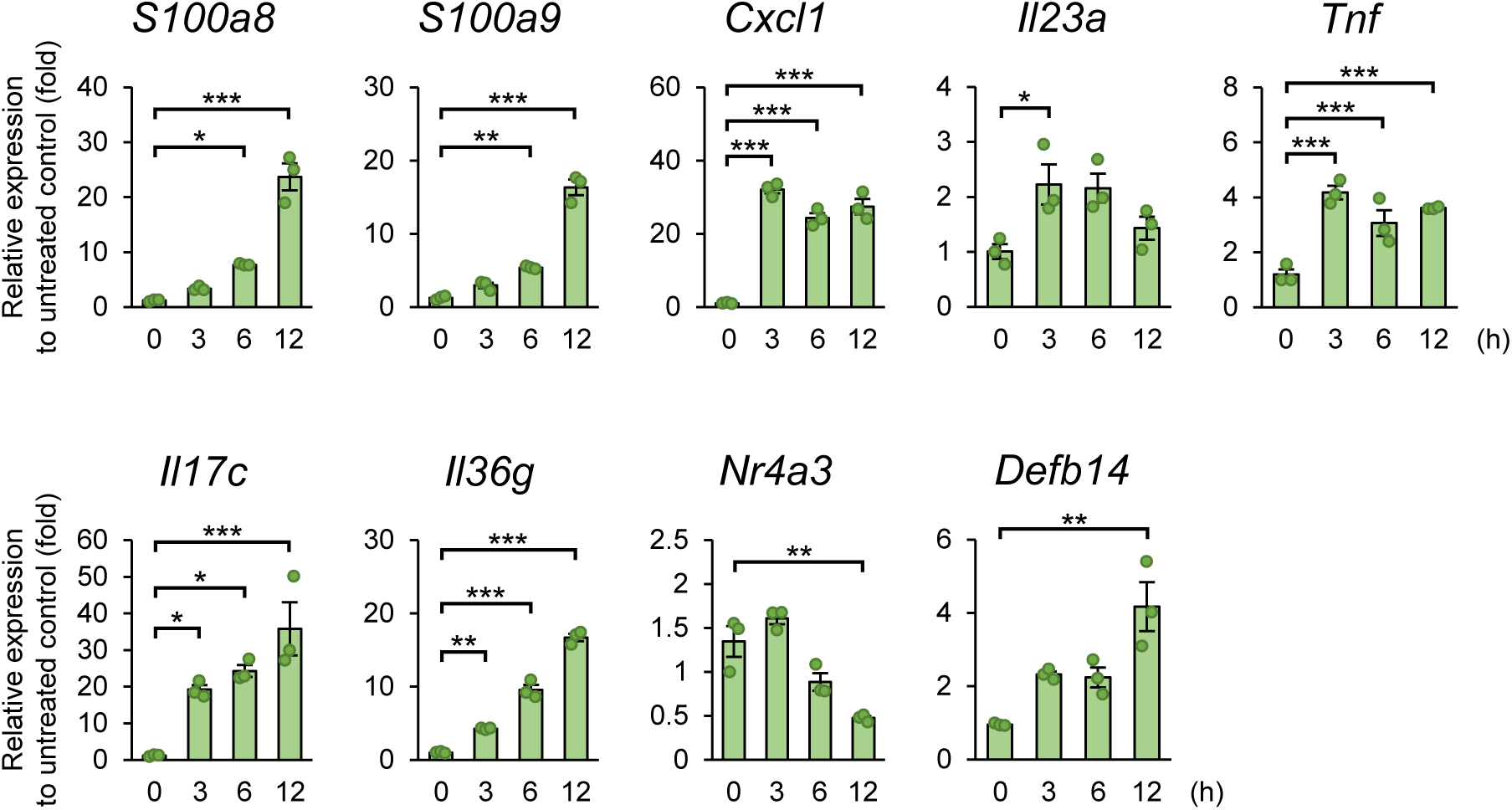
Expression of psoriasis-associated genes in primary mouse keratinocytes following IL-17A and TNF-α stimulation. Primary mouse keratinocytes were stimulated with IL-17A and TNF-α, and the expression of the indicated genes was analyzed by qPCR (*n* = 3). Data are presented as the mean ± s.e.m. Each circle indicates an independent biological sample. *P*-values were calculated by one-way ANOVA with Tukey’s test. (*: *p* < 0.05, **: *p* < 0.01, ***: *p* < 0.001).

**Supplemental Figure 7.**
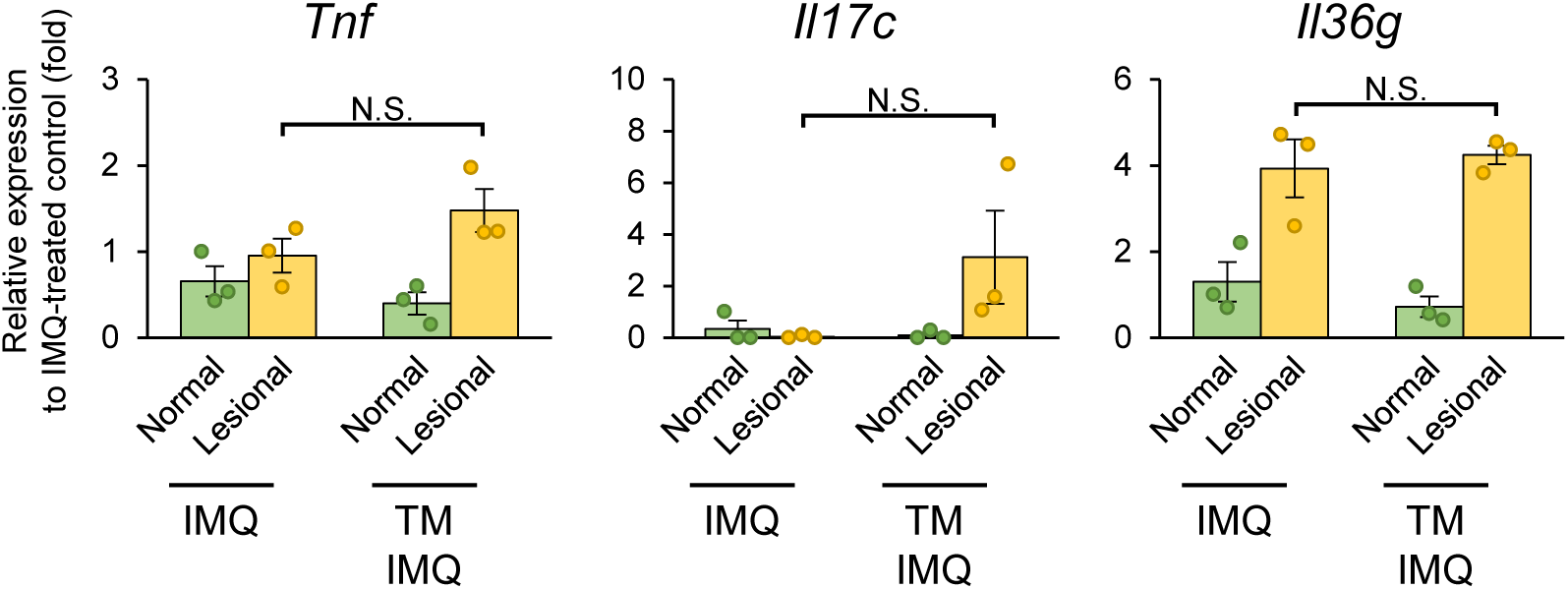
Tunicamycin does not alter the expression of Tnf, Il17c, or Il36g in IMQ-induced dermatitis. Mice were pretreated with tunicamycin (Tm) or vehicle prior to topical application of IMQ cream as described in Figure 4. The mRNA expression levels of *Tnf*, *Il17c*, and *Il36g* in ear tissues were determined by RT-qPCR (*n* = 3). Data are presented as the mean ± s.e.m. Each circle indicates an independent biological sample. *P*-values were calculated by one-way ANOVA with Tukey’s test. (N.S.: not significant).

**Supplemental Figure 8.**
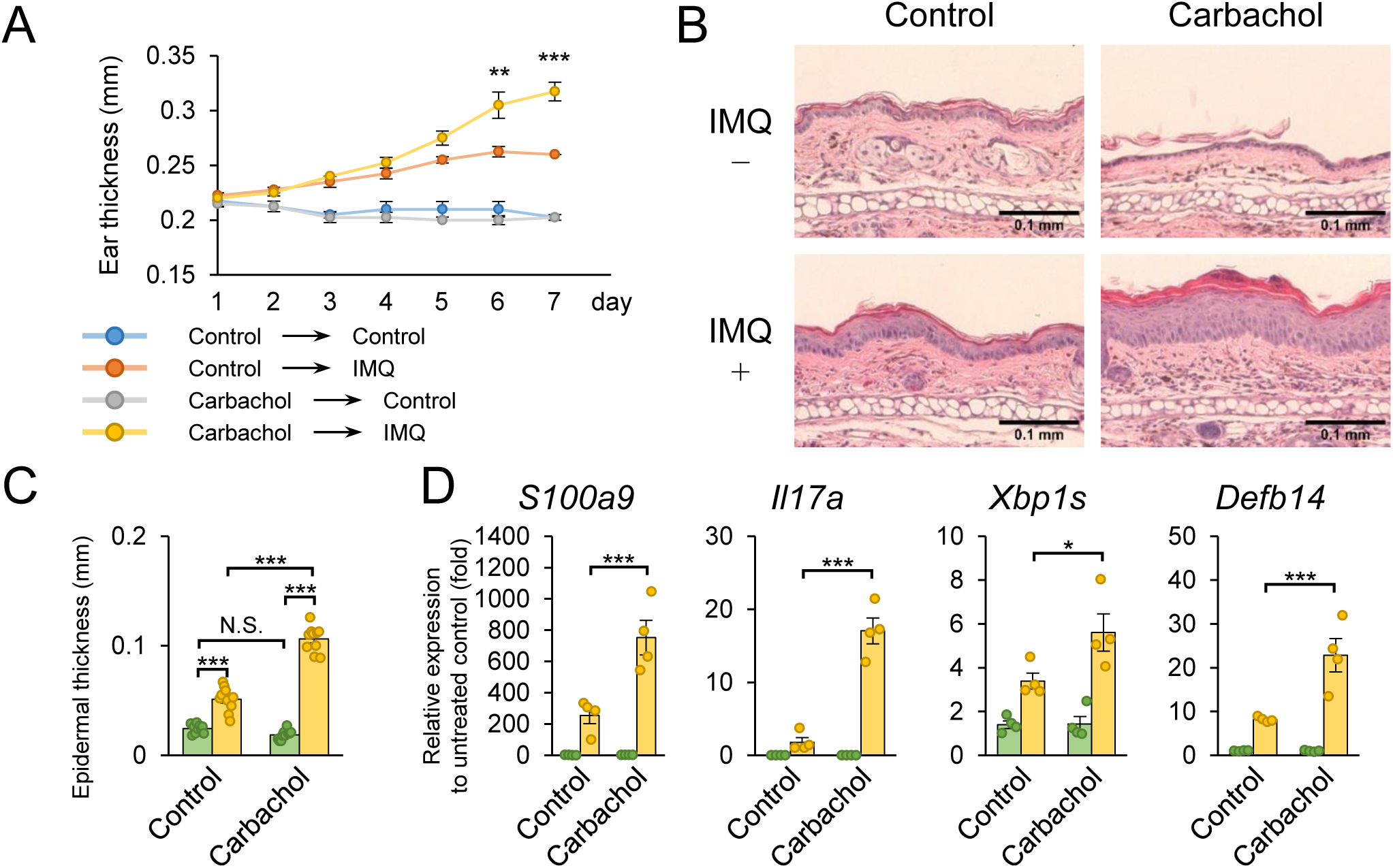
Carbachol treatment worsened IMQ-induced dermatitis. **(A-D)** IMQ cream was applied daily to the ear lobes of mice together with an intraperitoneal injection of carbachol. Ear thickness was measured daily (n = 4) (A). Mouse ears were sampled and subjected to histological analysis (H&E staining). Data are shown as representative histological analysis images (B). The epidermal thickness in (B) was measured (n = 10) (C). Total RNA was harvested from the ears and the expressions of S100a9, Il17a, Xbp1s, and Defb14 mRNA were measured by qPCR (n = 4) (D). Data are presented as the mean ± s.e.m. Each circle indicates an independent biological sample. *P*-values were calculated using one-way ANOVA with Tukey’s test. (N.S.: not significant, *: p < 0.05, **: p < 0.01, ***: p < 0.001).

**Supplemental Figure 9.**
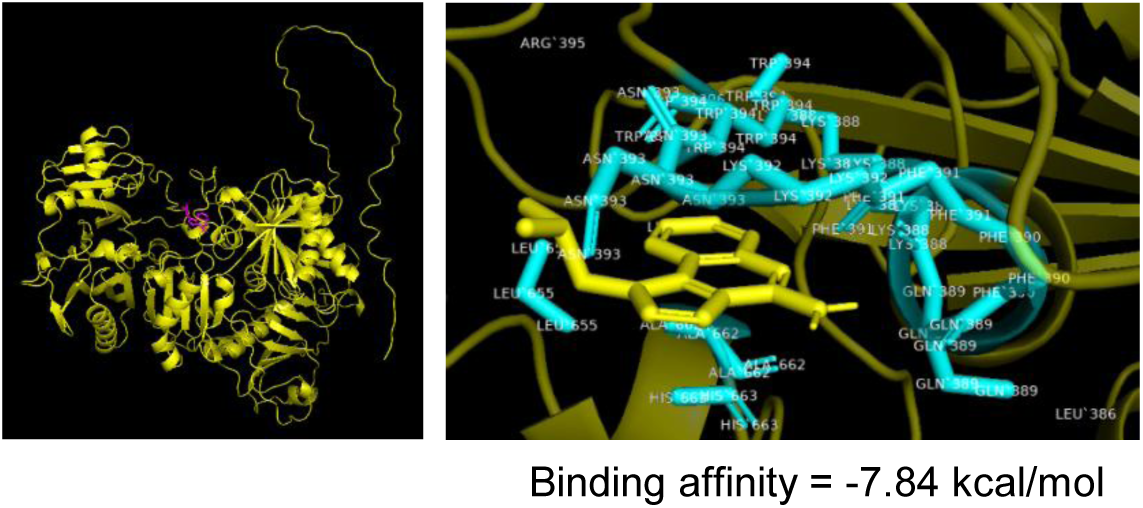
Surface plasmon resonance analysis between Gelsolin and IMQ. (Left panel) Yellow: Gelsolin, Perple: IMQ (Right panel) Right blue: amino acid residues of Gelsolin located near IMQ, Right yellow: IMQ

**Supplemental Figure 10.**
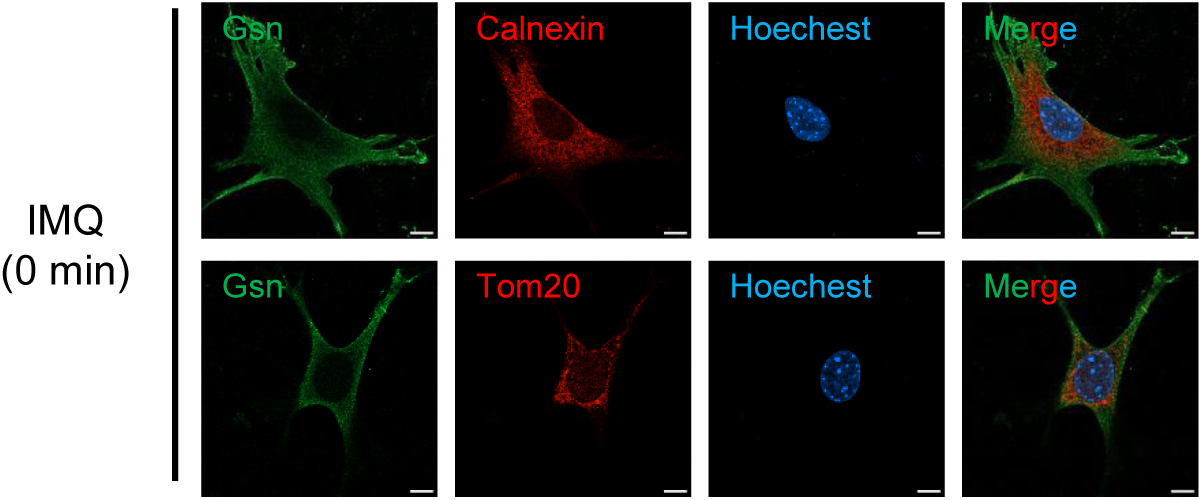
Subcellular localization of Gelsolin. FLAG-Gelsolin was transiently expressed in MEFs and cells were stimulated with IMQ. Gelsolin was visualized in green, Calnexin (ER) and TOM20 (mitochondria) were stained in red, and nuclei were counterstained with Hoechst. Data are representative images of two independent experiments.

**Supplemental Figure 11.**
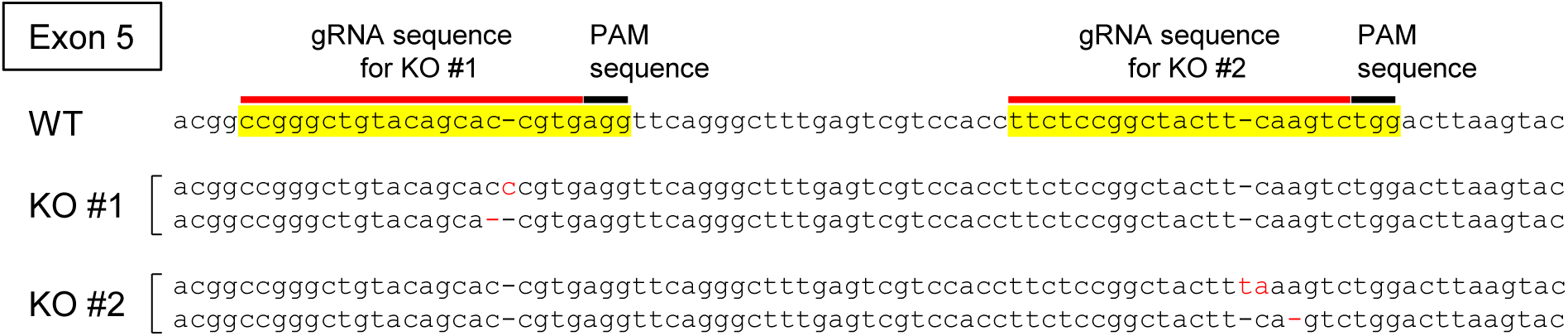
Genotyping of Gelsolin-deficient MEFs.

**Supplemental Figure 12.**
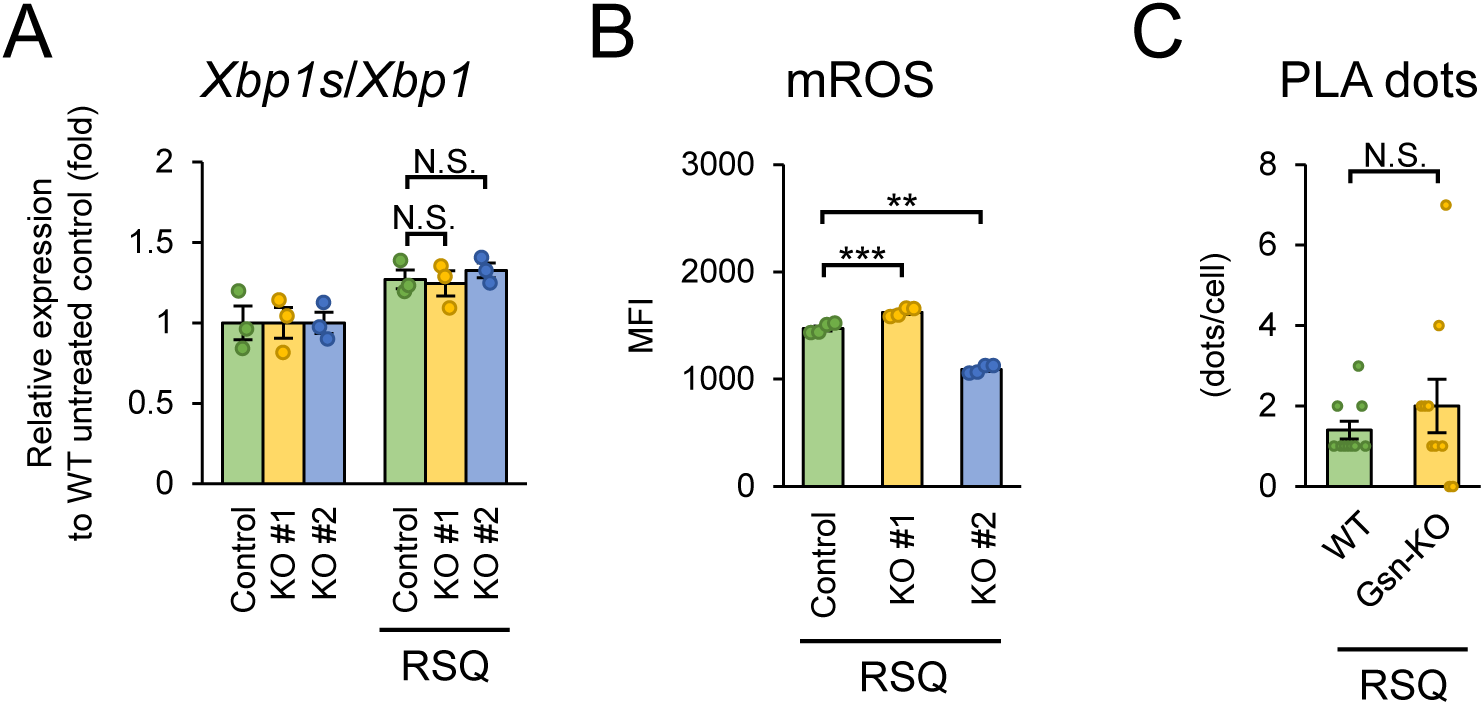
The effects of Gelsolin deficiency are specific to IMQ but not R848 stimulation. a, WT (Control) and Gsn-deficient MEFs were stimulated with RSQ, and Xbp1s expression was measured by qPCR analysis (*n* = 3). b, WT (Control) and Gsn-deficient MEFs were pre-stained with MitoSOX Red reagent, stimulated with RSQ, and mitochondrial ROS production was analyzed by flow cytometry (*n* = 4). c, WT (Control) and Gsn-deficient MEFs were pre-treated with LPS, followed by RSQ stimulation, and VDAC1–IP3R1 interactions were analyzed by PLA (*n* = 10). Data are presented as the mean ± s.e.m. Each circle indicates an independent biological sample. P-values were calculated by Student’s *t*-test. (N.S.: not significant, **: p < 0.01, ***: p < 0.001).

## Notes

### Competing Interest Statement

The authors have declared no competing interest.

### Summary of Updates

This revised version includes clarification of the interpretation of the IMQ-induced dermatitis model, additional keratinocyte experiments and comparative analyses, updated figures, methods, and supplementary files, and revised author affiliations.

